# Searching for rewards like a child means less generalization and more directed exploration

**DOI:** 10.1101/327593

**Authors:** Eric Schulz, Charley M. Wu, Azzurra Ruggeri, Björn Meder

## Abstract

How do children and adults differ in their search for rewards? We consider three different hypotheses that attribute developmental differences to either children’s increased random sampling, more directed exploration towards uncertain options, or narrower generalization. Using a search task in which noisy rewards are spatially correlated on a grid, we compare 55 younger children (age 7-8), 55 older children (age 9-11), and 50 adults (age 19-55) in their ability to successfully generalize about unobserved outcomes and balance the exploration-exploitation dilemma. Our results show that children explore more eagerly than adults, but obtain lower rewards. Building a predictive model of search to disentangle the unique contributions of the three hypotheses of developmental differences, we find robust and recoverable parameter estimates indicating that children generalize less and rely on directed exploration more than adults. We do not, however, find reliable differences in terms of random sampling.

## Introduction

Alan Turing (1950) famously believed that in order to build a General Artificial Intelligence, one must create a machine that can learn like a child. Indeed, recent advances in machine learning often contain references to child-like learning and exploration (Riedmiller et al., 2018). Yet little is known about how children actually explore and search for rewards in their environments, and in what ways their behavior differs from adults.

In the course of learning through interactions with the environment, all organisms (biological or machine) are confronted with the *exploration-exploitation dilemma* (Mehlhorn et al., 2015). This dilemma highlights two opposing goals. The first is to explore unfamiliar options that provide useful information for future decisions, yet may result in poor immediate rewards. The second is to exploit options known to have high expectations of reward, but potentially forgo learning about unexplored options.

In addition to balancing exploration and exploitation, another crucial ingredient for adaptive search behavior is a mechanism that can *generalize* beyond observed outcomes, thereby guiding search and decision making by forming inductive beliefs about novel options. For example, from a purely combinatorial perspective, it only takes a few features and a small range of values to generate a pool of options vastly exceeding what could ever be explored in a lifetime. Nonetheless, humans of all ages manage to generalize from limited experiences in order to choose from amongst a set of potentially unlimited possibilities. Thus, a model of human search also needs to provide a mechanism for generalization.

Previous research has found extensive variability and developmental differences in children’s and adults’ search behavior, which not only result from a progressive refinement of basic cognitive functions (e.g., memory or attention), but also derive from systematic changes in the computational principles driving behavior (Palminteri, Kilford, Coricelli, & Blakemore, 2016). In particular, developmental differences in learning and decision making have been explained by appealing to three hypothesized mechanisms: children sample more randomly, explore more eagerly, or generalize more narrowly than adults.

In this paper, we investigate how these three mechanisms are able to explain developmental differences in exploration-exploitation behavior. We provide a precise characterization of these competing ideas in a formal model, which is used to predict behavior in a search task, where noisy and continuous rewards are spatially correlated. Using behavioral markers, interpreting parameter estimates from computational models, and analyzing judgments about unexplored options, our results converge on the finding that children generalize less, but engage in more directed exploration than adults. We do not, however, find reliable developmental differences in random exploration. These results enrich our understanding of maturation in learning and decision making, demonstrating that children explore using uncertainty-guided mechanisms rather than simply behaving more randomly.

### A tale of three mechanisms

#### Development as cooling off

Because optimal solutions to the exploration-exploitation dilemma are generally intractable (Bellman, 1952), heuristic alternatives are frequently employed. In particular, learning under the demands of the exploration-exploitation trade-off has been described using at least two distinct strategies (Wilson, Geana, White, Ludvig, & Cohen, 2014). One such strategy is increased *random exploration*, which uses noisy, random sampling to learn about new options.

A key finding in the psychological literature is that children tend to try out more options than adults (Cauffman et al., 2010; Mata, Wilke, & Czienskowski, 2013). This has been interpreted as evidence for higher levels of random exploration in children, and has been loosely compared to algorithms of simulated annealing from computer science (Gopnik et al., 2017), where the amount of random exploration gradually reduces over time. Children can be described as having higher temperature parameters, where the learner initially samples very randomly across a large set of possibilities, before eventually focusing on a smaller subset (Gopnik, Griffiths, & Lucas, 2015). This temperature parameter is expected to “cool off” with age, leading to lower levels of random exploration in late childhood and adulthood.

#### Development as reduction of directed exploration

A second strategy to tackle the exploration-exploitation dilemma is to use *directed exploration* by preferentially sampling highly uncertain options in order to gain more information and reduce uncertainty about the environment. Directed exploration has been formalized by introducing an “uncertainty bonus” that values the exploration of lesser known options (Auer, 2002), with behavioral markers found in a number of studies (Frank, Doll, Oas-Terpstra, & Moreno, 2009; Wu, Schulz, Speekenbrink, Nelson, & Meder, 2018).

Directed exploration treats information as intrinsically valuable by inflating rewards by their estimated uncertainty (Auer, 2002). This leads to a more sophisticated *uncertainty-guided sampling* strategy that could also explain developmental differences. Indeed, the literature on self-directed learning shows that children are clearly capable of exploring their environment in a systematic, directed fashion. Already infants tend to value the exploration of uncertain options (L. E. Schulz, 2015), and children can balance theory and evidence in simple exploration tasks (Bonawitz, van Schijndel, Friel, & Schulz, 2012) and are able to efficiently adapt their search behavior to different environmental structures (Ruggeri & Lombrozo, 2015). Moreover, children can sometimes even outperform adults in the self-directed learning of unusual relationships (Lucas, Bridgers, Griffiths, & Gopnik, 2014). Both directed and random exploration do not have to be mutually exclusive mechanisms, with recent research finding signatures of both types of exploration in adolescent and adult participants (Gershman, 2018; Somerville et al., 2017; Wilson et al., 2014).

#### Development as refined generalization

Rather than explaining development as a change in how we explore given some beliefs about the world, *generalization-based accounts* attribute developmental differences to the way we form our beliefs in the first place. Many studies have shown that human learners use structured knowledge about the environment to guide exploration (E. Schulz, Konstantinidis, & Speekenbrink, 2017), where the quality of these representations and the way that people utilize them to generalize across experiences can have a crucial impact on search behavior. Thus, development of more complex cognitive processes (Blanco et al., 2016), leading to broader generalizations, could also account for the observed developmental differences in sampling behavior.

The notion of generalization as a mechanism for explaining developmental differences has a long standing history in psychology. For instance, Piaget (1964) assumed that children learn and adapt to different situational demands by the processes of assimilation (applying a previous concept to a new task) and accommodation (changing a previous concept in the face of new information). Expanding on Piaget’s idea, Klahr (1982) proposed generalization as a crucial developmental process, in particular the mechanism of regularity detection, which supports generalization and improves over the course of development. More generally, the implementation of various forms of decision making (Hartley & Somerville, 2015) could be constrained by the capacity for complex cognitive processes, which become more refined over the life span. For example, although younger children attend more frequently to irrelevant information than older children (Hagen & Hale, 1973), they can be prompted to attend to the relevant information by marking the most relevant cues, whereupon they eventually select the best alternative (Davidson, 1996). Thus, children may indeed be able to apply uncertainty-driven exploratory strategies, but lack the appropriate task representation to successfully implement them.

## A task to study generalization and exploration

We study the behavior of both children and adults in a *spatially correlated multi-armed bandit* task (Fig. 1a; Wu et al., 2018), where rewards are distributed on a grid characterized by spatial correlation (i.e., similar rewards cluster together; Fig. 1g; see also White, 2013, for a similar task) and the search horizon is vastly smaller than the number of options. Efficient search and accumulation of rewards in such an environment requires two critical components. First, participants need to learn about the underlying spatial correlation in order to generalize from observed rewards to unseen options. This is crucial because there are considerably more options than can be explored within the limited search horizon. Second, participants need a sampling strategy that achieves a balance between exploring new options and exploiting known options with high rewards.

**Figure 1.**
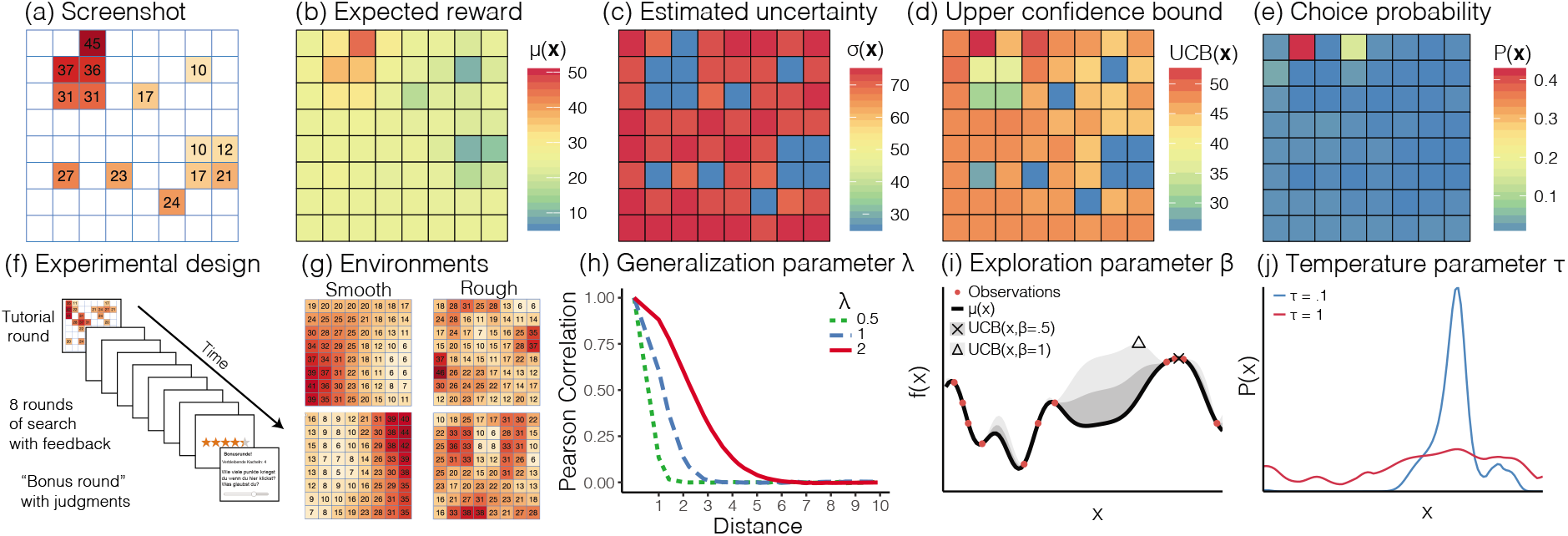
Overview of task and model. (a) Screenshot of experiment in the middle of a round with partially revealed grid. Expected reward (b) and estimated uncertainty (c) based on observations in (a) using Gaussian Process regression as a model of generalization. (d) Upper confidence bounds of each option based on a weighted sum of panels b and c. (e) Choice probabilities of softmax function. Panels (b-e) use median participant parameter estimates. (f) Overview of the experimental design (g) and types of environments. (h) Correlations of rewards between different options decay exponentially as a function of their distance, where higher values of λ lead to slower decays and broader generalizations. (i) An illustration of UCB sampling using a univariate example, where the expected reward (black line) and estimated uncertainty (gray ribbon; for different values of *β*) are summed up. Higher values of *β* value the exploration of uncertain options more strongly (compare the argmax of the two beta values, indicated by the cross and the triangle). (j) Overview of softmax function, where higher values of the temperature parameter *τ* lead to increased random exploration.

## Methods

### Participants

We recruited 55 younger children (range: 7 to 8, 26 female, *M_age_*=7.53; SD=0.50), 55 older children (range: 9 to 11, 24 female, *M_age_*=9.95; SD=0.80), and 50 adults (range: 18 to 55, 25 female, *M_age_*=33.76; SD= 8.53) from the Berlin Natural History Museum in Germany. We determined the different age groups and the number of participants per group before data collection commenced, based on existing findings showing strong developmental differences between age 7 and 10 in children’s question-asking and active search behavior (Davidson, 1991; Ruggeri & Lombrozo, 2015). Participants were paid up to €3.50 for taking part in the experiment, contingent on performance (range: €2.00 to €3.50, *M_reward_*=€2.67; SD=0.50). Informed consent was obtained from all participants.

### Design

The experiment used a between-subjects design, where participants were randomly assigned to one of two different classes of environments (Fig. 1g), with *smooth environments* having stronger spatial correlations than *rough environments.* We generated 40 of each class of environments from a radial basis function kernel (see below), with *λ_smooth_* = 4 and *λ_rough_* = 1. On each round, a new environment was sampled (without replacement) from the set of 40 environments, which was then used to define a bivariate function on the grid, with each observation including additional normally distributed noise 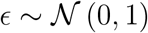. The task was presented over ten rounds on different grid worlds drawn from the same class of environments. The first round was a tutorial round and the last round was a bonus round in which participants sampled for 15 trials and then had to generate predictions for five randomly chosen and previously unobserved tiles on the grid. Participants had a search horizon of 25 trials per grid, including repeat clicks.

### Materials and procedure

Participants were introduced to the task through a tutorial round, which familiarized them with the spatial correlation of rewards and the possibility of re-clicking tiles. Moreover, participants were told that they would be rewarded based on the sum of sampled points. Afterwards, they had to complete three comprehension questions before starting the task. At the beginning of each round, one random tile was revealed and participants could click on any of the tiles (including re-clicks) on the grid until the search horizon was exhausted. Clicking an unrevealed tile displayed the numerical value of the reward along with a corresponding color aid, where darker colors indicated higher rewards. Per round, observations were scaled to a randomly drawn maximum value in the range of 35 to 45, so that the value of the global optima could not be easily guessed. Re-clicked tiles could show some variations in the observed value due to noise. For repeat clicks, the most recent observation was displayed numerically, while the color of the tile corresponded to the mean of all previous observations. In the bonus round, participants sampled for 15 trials and were then asked to generate predictions for five randomly selected and previously unobserved tiles. This was explained to them before the bonus round started. Additionally, participants had to indicate how certain they were about each prediction on a scale from 0 to 10. Afterwards, they had to select one of the five tiles before continuing with the round.

Participants were awarded up to five stars at the end of each round (e.g., 4.6 out of 5), based on the ratio of their average reward to the global maximum. The performance bonus was calculated based on the average number of stars earned in each round, excluding the tutorial round. 5 out of 5 stars corresponded to €3.50, while each half star interval reduced the bonus by €0.50 until a minimum bonus of €0.50.

### A combined model of generalization and exploration

We use a formal model that combines generalization with a sampling strategy accounting for both directed and random exploration (Wu et al., 2018), and use it to predict each participant’s out-of-sample search behavior. The generalization component is based on *Gaussian Process* (GP) regression, which is a Bayesian function learning approach theoretically capable of learning any stationary function (Rasmussen & Williams, 2006) and has been found to effectively describe human behavior in explicit function learning tasks (Lucas, Griffiths, Williams, & Kalish, 2015). The GP component is used to adaptively learn a value function, which generalizes the limited set of observed rewards over the entire search space using Bayesian inference.

The GP prior is completely determined by the choice of a kernel function *k*(x, x′), which encodes assumptions about how points in the input space are related to each other. A common choice of this function is the *radial basis function*:

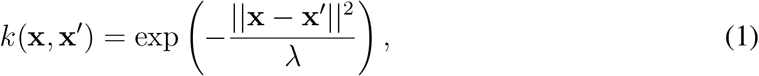

where the length-scale parameter λ encodes the extent of spatial generalization between options (tiles) in the grid. The assumptions of this kernel function are similar to the gradient of generalization historically described by Shepard (1987), which also models generalization as an exponentially decaying function of the stimulus similarity distance (see Fig. 1h), which has been observed across a wide range of stimuli and organisms. As an example, generalization with λ =1 corresponds to the assumption that the rewards of two neighboring tiles are correlated by *r* = 0.6, and that this correlation effectively decays to zero for options further than three tiles apart. We treat λ as a free parameter in our model comparison in order to assess age-related differences in the capacity for generalization.

Given different possible options x to sample from (i.e., tiles on the grid), GP regression generates normally distributed beliefs about rewards with expectation *μ*(x) and estimated uncertainty *σ*(x) (Fig. 1b,c). A sampling strategy is then used to map the beliefs of the GP onto a valuation for sampling each option at a given time. Crucially, such a sampling strategy must address the exploration-exploitation dilemma. One frequently applied heuristic for solving this dilemma is *Upper Confidence Bound* (UCB) sampling (Srinivas, Krause, Kakade, & Seeger, 2009), which evaluates each option based on a weighted sum of expected reward and estimated uncertainty:

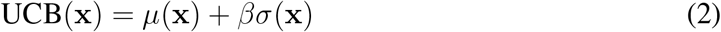

where *β* models the extent to which uncertainty (in addition to mean rewards) is valued positively and therefore directly sought out. This strategy corresponds to directed exploration because it encourages the sampling of options with higher uncertainty according to the underlying generalization model (see Fig. 1i). We treat the exploration parameter *β* as a free parameter to assess how much participants value the reduction of uncertainty (i.e., engage in directed exploration). As an example, an exploration bonus of *β* = 0.5 means participants would prefer an option *x*_1_ expected to have reward *μ*(*x*_1_) = 30 and uncertainty *σ*(*x*_1_) = 10, over option *x*_2_ expected to have reward *μ*(*x*_2_) = 34 and uncertainty *σ*(*x*_2_) = 1. This is because sampling *x*_1_ is expected to reduce a larger amount of uncertainty, even though *x*_2_ has a higher expected reward (UCB(*x*_1_|*β* = 0.5) = 35 vs. UCB(*x*_2_|*β* = 0.5) = 34.5).

Finally, we use a softmax function to map the upper confidence bound values, UCB(x), of our proposed Gaussian Process-Upper Confidence Bound sampling model onto choice probabilities:

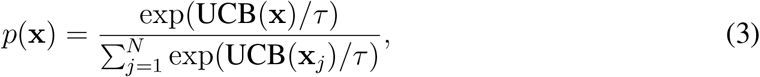

where *τ* is the temperature parameter governing the amount of randomness in sampling behavior. If *τ* is high (higher temperatures), then participants are assumed to sample more randomly, whereas if *τ* is low (cooler temperatures), the choice probabilities are concentrated on the highest valued options (Fig. 1j). Thus, *τ* encodes the tendency towards random exploration. We treat *τ* as a free parameter to assess the extent of random exploration in children and adults (see Supplemental Material for alternative implementations such as *∊*-greedy sampling and estimation of optimal parameters).

In summary, GP-UCB contains three different parameters: the length-scale *λ* capturing the extent of generalization, the exploration bonus *β* describing the extent of directed exploration, and the temperature parameter *τ* modulating random exploration. These three parameters directly correspond to the three postulated mechanisms of developmental differences in various decision making tasks and can also be robustly recovered (see Supplemental Material).

## Results

### Behavioral results

Participants gained higher rewards in smooth than in rough environments (Fig. 2a; comparing participants’ average rewards: *t*(158) = 10.51, *p* < .001, *d* = 1.66, 95% CI=[1.30,2.02], *BF* > 100), suggesting they made use of the spatial correlations and performed better when correlations were stronger. Adults performed better than older children (Fig. 2a; *t*(103) = 4.91, *p* < .001, *d* = 0.96, 95% CI=[0.55,1.37], *BF* > 100), who in turn performed somewhat better than younger children (*t*(108) = 2.42, *p* = .02, *d* = 0.46, 95% CI=[0.08, 0.84], *BF* = 2.68). Analyzing the distance between consecutive choices (Fig. 2b) revealed that participants sampled more locally (smaller distances) in smooth compared to rough environments (*t*(158) = −3.83, *p* < .001, *d* = 0.61, 95% CI=[0.29, 0.93], *BF* > 100). Adults sampled more locally than older children (*t*(103) = −3.9, *p* < .001, *d* = 0.76, 95% CI=[0.36,1.16], *BF* > 100), but there was no difference between younger and older children (*t*(108) = 1.76, *p* = .08, *d* = 0.34, 95% CI=[−0.05, 0.72], *BF* = 0.80). Importantly, adults sampled fewer unique options than older children (14.5 vs. 21.7; *t*(103) = 6.77, *d* = 1.32, 95% CI=[0.90,1.75], *p* < .001, *BF* > 100), whereas the two children groups did not differ in how many unique options they sampled (21.7 vs. 22.7; *t*(108) = 1.27, *d* = 0.24, 95% CI=[−0.14, 0.62], *p* = .21, *BF* = 0.4).

**Figure 2.**
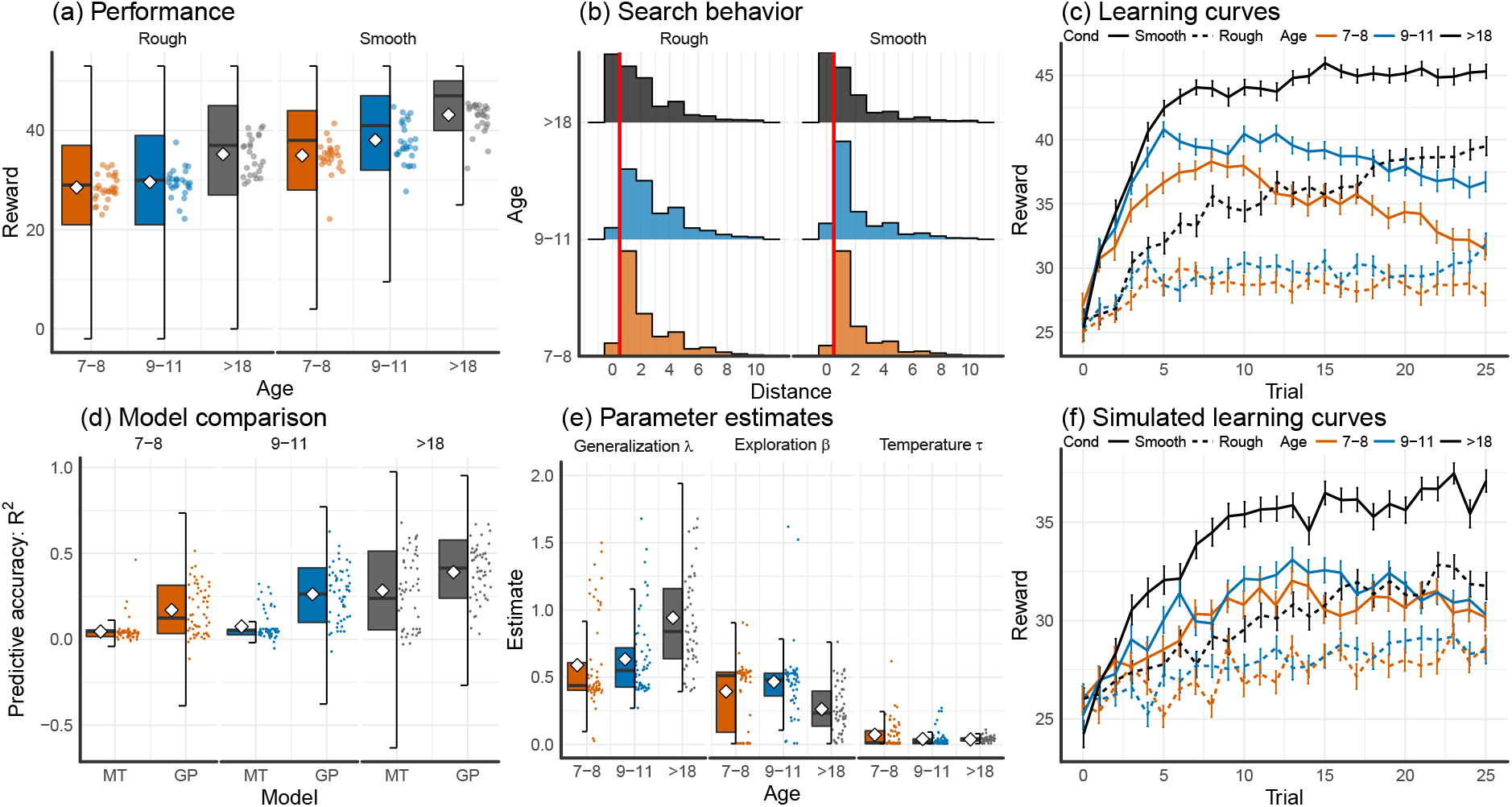
Main results. (a) Tukey box plots of rewards, showing the distribution of all choices for all participants, with the horizontal line representing the median and box showing the interquartile range of the distribution. Each dot is the participant-wise mean and diamonds indicate group means. (b) Histograms of distances between consecutive choices by age group and condition, with a distance of zero corresponding to a repeat click. The vertical red line marks the difference between a repeat click and sampling a different option. (c) Mean reward over trials by condition (solid lines for smooth and dashed lines for rough environments) and age group (color). Error bars indicate the standard error of the mean. (d) Tukey box plots showing the results of the model comparison between Gaussian Process (GP) and Mean Tracker (MT) models by age group. Each point is a single subject and group means are shown as a diamond. (e) Tukey box plot of cross-validated parameters retrieved from the GP-UCB model by age group, where each point is the mean estimate per subject and diamonds indicate the group means. Outliers are removed for readability, but are included in all statistical tests (see Supplemental Material). (f) Learning curves simulated by GP-UCB model using mean participant parameter estimates. Error bars indicate the standard error of the mean.

Looking at the learning curves (i.e., average rewards over trials; Fig. 2c), we found a positive rank-correlation between mean rewards and trial number (Spearman’s *ρ* = .12, *t*(159) = 6.12, *p*< .001, 95% CI=[0.08, 0.16], *BF* > 100). Although this correlation did not differ between the rough and smooth condition (*t*(158) = −0.43, *p* = .67, *d* = 0.07, 95% CI=[−0.24, 0.38], *BF* = 0.19), it was significantly higher for adults than for older children (0.29 vs 0.08, *t*(103) = 5.90, *p*< .001, *d* = 1.15, 95% CI=[0.74,1.57], *BF* = 0.19, *BF* > 100). The correlation between trials and rewards did not differ between younger and older children (0.04 vs 0.08; *t*(108) = −1.87, *p* = .06, *d* = 0.36, 95% CI=[−0.02,0.74], *BF* = 0.96). Therefore, adults learned faster, while children explored more extensively (see Supplemental Material for further behavioral analyses).

### Model comparison

We compared the GP-UCB model with an alternative model that does not generalize across options but is a powerful Bayesian model for reinforcement learning across independent reward distributions *(Mean Tracker*; MT). Model comparisons are based on leave-one-round-out cross-validation error, where we fit each model combined with the UCB sampling strategy to each participant using a training set omitting one round, and then assess predictive performance on the hold-out round. Repeating this procedure for every participant and all rounds (apart from the tutorial and the bonus rounds), we calculated the standardized *predictive accuracy* for each model (pseudo-*R*^2^ comparing out-of-sample log loss to random chance), where 0 indicates chance-level predictions and 1 indicates theoretically perfect predictions (see Supplemental Material for full model comparison with additional sampling strategies). The results of this comparison are shown in Fig. 2d. The GP-UCB model predicted participants’ behavior better overall (*t*(159) = 13.28, *p* < .001, *d* = 1.05, 95% CI=[0.82,1.28], *BF* > 100), and also for adults (*t*(49) = 5.98, *p* < .001, *d* = 0.85, 95% CI=[0.43,1.26], *BF* > 100), older (*t*(54) = 10.92, *p* < .001, *d* = 1.48, 95% CI=[1.05,1.90], *BF* > 100) and younger children (*t*(54) = 6.77, *p* < .001, *d* = 0.91, 95% CI=[0.52,1.31], *BF* > 100). The GP-UCB model predicted adults’ behavior better than that of older children (*t*(103) = 4.33, *p* < .001, *d* = 0.85, 95% CI=[0.44,1.25], *BF* > 100), which in turn was better predicted than behavior of younger children (*t*(108) = 3.32, *p* = .001, *d* = 0.63, 95% CI=[0.24,1.02], *BF* = 24.8).

### Developmental differences in parameter estimates

We analyzed the mean participant parameter estimates of the GP-UCB model (Fig. 2e) to assess the contributions of the three mechanisms (generalization, directed exploration, and random exploration) towards developmental differences.

We found that adults generalized more than older children, as indicated by larger λ-estimates (Mann-Whitney-*U* = 2001, *p*< .001, *r_τ_* = 0.32, 95% CI=[0.18, 0.47], *BF* > 100), whereas the two groups of children did not differ significantly in their extent of generalization (*U* = 1829, *p* = .06, *r_τ_* = 0.15, 95% CI=[−0.01, 0.30], *BF* = 1.7). Furthermore, older children valued the reduction of uncertainty more than adults (i.e., higher β-values; *U* = 629, *p* < .001, *r_τ_* = 0.39, 95% CI=[0.25, 0.52], *BF* > 100), whereas there was no difference between younger and older children (*U* = 1403, *p* = .51, *r_τ_* = 0.05, 95% CI=[−0.10, 0.21], *BF* = 0.2). Critically, whereas there were strong differences between children and adults for the parameters capturing generalization and directed exploration, there was no reliable difference in the softmax temperature parameter *τ*, with no difference between older children and adults (*W* = 1718, *p* = .03, *r_τ_* = 0.17, 95% CI=[0.01,0.34], *BF* = 0.7) and only anecdotal differences between the two groups of children (*W* = 1211, *p* = .07, *r_τ_* = 0.14, 95% CI=[−0.01, 0.30] *BF* = 1.4).^1^ This suggests that the amount of random exploration did not reliably differ by age group (see Supplemental Materials for other implementations of random exploration). Thus, our modeling results converge on the same conclusion as the behavioral results. Children explore more than adults, yet instead of exploring randomly, children’s exploration behavior seems to be directed toward options with high uncertainty. Additionally, our parameter estimates are robustly recoverable (see Supplemental materials) and can be used to simulate learning curves that reproduce the differences between the age groups as well as between smooth and rough conditions (Fig. 2f).

### Bonus round

In the bonus round, each participant predicted the expected rewards and the underlying uncertainty for five randomly sampled unrevealed tiles after having made 15 choices on the grid. We first calculated the mean absolute error between predictions and the true expected value of rewards (Fig. 3a). Prediction error was higher for rough compared to smooth environments (*t*(158) = 4.93, *p* < .001, *d* = 0.78, 95% CI= [0.46,1.10], *BF* > 100), reflecting the lower degree of spatial correlation that could be utilized to evaluate unseen options. Surprisingly, older children were as accurate as adults (*t*(103) = 0.28, *p* = .78, *d* = 0.05, 95% CI=[−0.44,0.33], *BF* = 0.2), but younger children performed worse than older children (*t*(108) = 3.14, *p* = .002, *d* = 0.60, 95% CI=[0.21,0.99], *BF* = 15). Certainty judgments did not differ between the smooth and rough environments (*t* (158) = 1.13, *p* = .26, *d* = 0.18, 95% CI=[−0.13, 0.49], *BF* = 0.2) nor between the different age groups (max-*BF* = 0.1).

**Figure 3.**
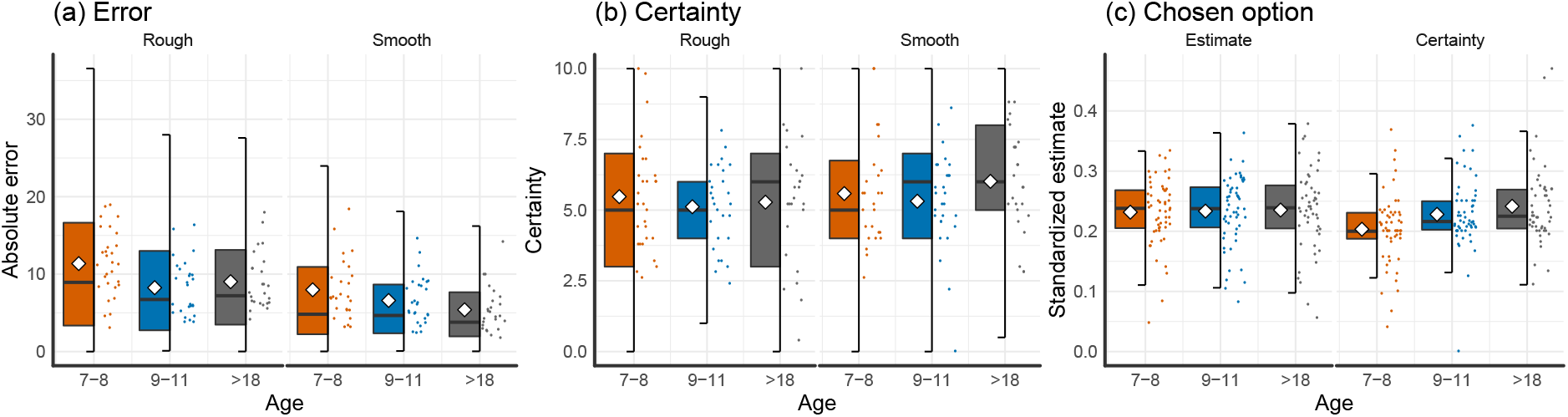
Bonus round results. (a) Mean absolute error of participant predictions about the rewards of unobserved tiles. (b) Certainty judgments, where 0 is least certain and 10 is most certain. (c) Standardized predictions and certainty estimates, which shows how much the estimated reward and certainty influenced choice (relative to judgments about non-chosen options). All figures show Tukey box plots (over all data points), with participant means as dots and group means as diamonds.

Of particular interest is how judgments about the expectation of rewards and perceived uncertainty relate to the eventual choice from amongst the five options (implemented as a 5-alternative forced choice). We standardized the estimated reward and confidence judgment of each participant’s chosen tile by dividing by the sum of the estimates for all five options (Fig. 3c). Thus, larger standardized estimates reflect a larger contribution of either high reward or high certainty on the choice. Whereas there was no difference between age groups in terms of the estimated reward of the chosen option (max-*BF* = 0.1), we found that younger children preferred options with higher uncertainty slightly more than older children (*t*(108) = 2.22, *p* = .03, *d* = 0.42, 95% CI=[0.04,0.80], *BF* = 1.8), and substantially more than adults (*t*(103) = 2.82, *p* = .006, *d* = 0.55, 95% CI=[0.16, 0.95], *BF* = 6.7). This further corroborates our previous analyses, showing that the sampling behavior of children is more directed toward uncertain options than that of adults.

## Discussion

We examined three potential sources of developmental differences in a complex learning and decision-making task: random exploration, directed exploration, and generalization. Using a paradigm that combines both generalization and search, we found that adults gained higher rewards and exploited more strongly, whereas children sampled more unique options, thereby gaining lower rewards but exploring the environment more extensively. Using a computational model with parameters directly corresponding to the three hypothesized mechanisms of developmental differences, we found that children generalized less and were guided by directed exploration more strongly than adults. They did not, however, explore more randomly than adults.

Our results shed new light on the developmental trajectories in generalization and exploration, casting children not as merely prone to more random sampling behavior, but as directed explorers who are hungry for information in their environment. Our conclusions are drawn from converging evidence combining analysis of behavioral data and computational modeling. Moreover, our findings are highly recoverable and also hold for other formalizations of random exploration instead of using the softmax temperature parameter (see Supplemental Materials).

Interestingly, related work by Somerville et al. (2017) also found no developmental difference in random exploration, but increasing directed exploration across early adolescence, which stabilized in adulthood. We believe that our results are not necessarily incompatible with that finding. Somerville and colleagues defined directed exploration using horizon-sensitive exploration (i.e,. strategic planning of exploration), whereas we define directed exploration as uncertainty-guided exploration via a greedy upper confidence bound algorithm. Thus, children may have higher tendencies towards directed exploration in a stepwise-greedy fashion, but fail to exhibit such tendencies when planning ahead for multiple steps, perhaps due to cognitive limitations. This opens up further possibilities for studying different mechanisms of directed exploration and how they relate to one another.

Our results provide strong evidence for developmental differences in directed exploration driven by both expected rewards and the associated uncertainty. These findings complement existing research on age-related differences in risk- and uncertainty-related behavior (Josef et al., 2016). For instance, adolescents and adults systematically differ in their tolerance of options with outcomes that have unknown probabilities, providing converging evidence that uncertainty is valued differently depending on age (Tymula et al., 2012). Importantly, in our task a sampling strategy that only seeks to reduce uncertainty is inferior to the “optimistic” UCB strategy in predicting children’s and adults’ behavior (see SOM-U for details). This result demonstrates how reward expectations and uncertainty interact to produce decision-making behavior that balances the exploration-exploitation trade-off adaptively as a function of age. Future work should attempt to further disentangle different interpretations of uncertainty seeking formally, for example, by not familiarizing participants with the underlying environments or by manipulating the level of noise in the outcomes directly.

Furthermore, it is surprising that there were no meaningful differences between younger and older children’s parameter estimates. Since this indicates that directed exploration might be present even earlier than expected, future studies could apply our paradigm to investigate exploration behavior in even younger children.

Our results showing a developmental increase in generalization can also be related to previous findings showing a developmental increase in the use of task structure knowledge in model-based reward learning (Decker, Otto, Daw, & Hartley, 2016). Because the generalization parameter *λ* can be mathematically equated to the speed of learning about the underlying function (Sollich, 1999), generalization and learning are inextricably linked in our task. There are however other uses of the term “generalization” in the psychological literature. For example, children are known to generalize words or categories more broadly, a tendency that decreases over time, trading-off with the capacity to form more precise episodic memories (Keresztes et al., 2017). While we focus on generalization in the sense used by Shepard (i.e., generalization across stimuli), it is an outstanding question how this type of generalization relates to word and category generalization. It would be a fruitful avenue for future research to connect these two domains in a unifying theory of generalization.

In our current study, we have assessed only environments with stationary reward distributions. However, given that children displayed increased exploration behavior, we believe that they could perform especially well in environments that change over rounds. Whether or not children would outperform adults in changing environments remains an important question for future research.

Ultimately, our results suggest that to fulfill Alan Turing’s dream of creating a child-like AI, we need to incorporate generalization and curiosity-driven exploration mechanisms (Riedmiller et al., 2018).

## Author Contributions

All authors developed the study concept and contributed to the study design. E. Schulz and C.M. Wu performed the data analysis and interpretation under the supervision of B. Meder and A. Ruggeri. E. Schulz and C.M. Wu drafted the manuscript, and B. Meder and A. Ruggeri provided critical revisions. All authors approved the final version of the manuscript for submission.

## Acknowledgements

We thank all of the families who participated in this research, the Berlin Natural History Museum where we conducted the study, Andreas Sommer for collecting the data, and Federico Meini for help with programming the experiment.

## Declaration of Conflicting Interests

The authors declared that they had no conflicts of interest with respect to their authorship or the publication of this article.

## Funding

This work was supported by the Max Planck Society and DFG-grant ME 3717/2-2 to BM. ES is supported by the Harvard Data Science Initiative. CMW is supported by the International Max Planck Research School on Adapting Behavior in a Fundamentally Uncertain World.

## Open Practices

All code and data have been made publicly available and can be accessed at https://git.io/vppsK

## Reviewed Supplementary Materials

### Statistical tests

We report both frequentist and Bayesian statistics throughout the paper. Whereas frequentist tests are reported as either Student’s *t*-tests (for the behavioral data and model comparisons) or Mann-Whitney-*U* tests (for parameter comparisons), we rely on Bayes factors (*BF*) to quantify the relative evidence the data provide in favor of the alternative hypothesis (*H_A_*) over the null (*H*_0_).

For testing hypotheses regarding the behavioral data and the model comparison, we use the default two-sided Bayesian *t*-test for independent samples using a Jeffreys-Zellner-Siow prior with its scale set to 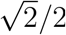, as suggested by Rouder, Speckman, Sun, Morey, and Iverson (2009). The prior is truncated below 0 for the directional tests performed to create Figure S1, which shows pairwise comparisons between the different models. All other statistical tests are non-directional as defined by a symmetric prior.

For testing hypotheses regarding the model parameters, we use the frequentist Mann-Whitney-*U* test and report Kendall’s *r_τ_* as an effect size. The Bayesian test is based on performing posterior inference over the test statistics and assigning a prior by means of a parametric yoking procedure (van Doorn, Ly, Marsman, & Wagenmakers, 2017). This then leads to a posterior distribution for Kendall’s *r_τ_*, and via the Savage-Dickey density ratio test, also yields an interpretable Bayes factor. The null hypothesis posits that parameters do not differ between the two groups, while the alternative hypothesis posits an effect and assigns an effect size using a Cauchy distribution with the scale parameter set to 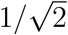.

We also report 95%-Confidence Intervals (95% CI) for both effect sizes, Cohen’s *d* (estimated directly) and *r_τ_* (bootstrapped estimators).

### Other forms of random exploration

A softmax function with a temperature parameter *τ* is only one way to define random exploration. Another approach towards assessing random exploration is so-called *ϵ*-greedy exploration. Given *k* number of arms (64 in our experiment), *ϵ*-greedy exploration chooses

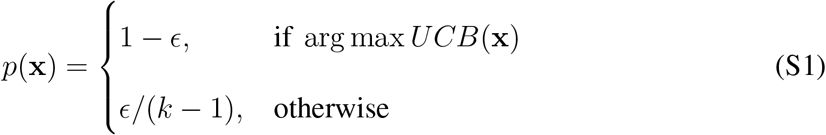

where *ϵ* is a free parameter. We test the *ϵ*-greedy method of exploration by using it instead of a softmax function in combination with the GP regression model and a UCB-sampling strategy. The results of this comparison show that the *ϵ*-greedy exploration model was systematically worse at predicting behavior than the softmax model reported in the main text (mean predictive accuracy: *R*^2^ = 0.21, *t*(159) = 6.67, *p* < .001, *d* = 0.53, 95% CI=[0.30, 0.75], *BF* > 100). Additionally, the softmax model also had better predictive accuracy than the *ϵ*-greedy exploration model for adults (mean predictive accuracy: *R*^2^ = 0.26, *t*(49) = 9.29 *p* < .001, *d* = 1.31, 95% CI=[0.88,1.75], *BF* > 100), and for older children (mean predictive accuracy: *R*^2^ = 0.21, *t*(54) = 3.60, *p* < .001, *d* = 0.49, 95% CI=[0.10, 0.87], *BF* = 38.5), but not for younger children (mean predictive accuracy: *R*^2^ = 0.17, *t*(54) = 0.33, *p* = .74, *d* = 0.04, 95% CI=[−0.33,0.42], *BF* = 0.2).

Next, we looked for age-related differences in the parameter estimates of the *ϵ*-greedy model^2^, specifically the directed exploration parameter *β* and the alternative random exploration parameter *ϵ*. As in the softmax-parameterized models, we find larger λ-estimates for adults than for older children (0.99 vs. 0.24, *U* = 1975, *r_τ_* = 0.31, 95% CI=[0.14, 0.46], *p* < .001, *BF* > 100), whereas the two children groups do not differ in their λ-estimates (0.31 vs. 0.24, *U* = 1299, *r_τ_* = 0.10, 95% CI=[−0.07, 0.24], *p* = .20, *BF* = 0.4). Furthermore, we find more directed exploration (larger *β* parameters) for older children than for adults (17.30 vs. 5.38, *U* = 555, *r_τ_* = 0.42, 95% CI=[0.30, 0.56], *p* < .001, *BF* > 100), but no difference between the two groups of children (17.20 vs. 17.30, *U* = 1684, *r_τ_* = 0.12, 95% CI=[−0.07, 0.24], *p* = .3, *BF* = 0.27). We also found a difference in *ϵ*-greedy exploration parameter between adults and older children (0.00012 vs. 0.00014, *U* = 960,*r_τ_* = 0.21, 95% CI=[0.07, 0.37], *p* = .007, *BF* = 6.95), but not between the two groups of children (0.00014 vs. 0.00016, *U* = 1774, *r_τ_* = 0.12, 95% CI=[−0.03, 0.28], *p* = .11, *BF* = 0.46). Notice that the relative proportion of random exploration decisions according to the *ϵ*-parameter estimates is so small, that over the 200 choices in our task, this accounts for a difference of approximately 1 in every 250 choices. Thus, there is almost no practical difference in participants’ e-parameters. The overall age-related effect in the *ϵ*-greedy analysis was also larger for directed exploration than for e-greedy exploration (*r_τ_* = 0.40 vs. *r_τ_* = 0.25).

Thus, there are two reasons to believe that children are driven more strongly by directed than by random exploration. Firstly, the GP-UCB model combined with a softmax formulation of random exploration predicted participants better than an *ϵ*-greedy model, and finds no age-related difference in terms of random exploration described by the temperature parameter *τ*. Secondly, parameter estimates of the e-greedy model find only small and practically meaningless age-related difference in *ϵ*-exploration, but again a large age-related differences in the directed exploration parameter *β*.

## Unreviewed Supplementary Materials

### Full modeling results

We report a full model comparison of two models of learning each combined with three different sampling strategies (see also Fig. S8S9). Different *models of learning* (i.e., Gaussian Process regression and Mean Tracker) are combined with different *sampling strategies*, in order to make predictions about where a participant will search next, given the history of previous observations. Table S1 contains the predictive accuracy, the number of participants best described, the log-loss, the probability of exceedance and the parameter estimates of each combination of a learning model and a sampling strategy. In total, cross-validated model comparisons for both models required approximately two days of computation time distributed on a cluster of 160 Dell C6220 nodes.

#### Models of Learning

##### Gaussian Process

We use Gaussian Process (GP) regression as a Bayesian model of generalization. A GP is defined as a collection of points, any subset of which is multivariate Gaussian. Let 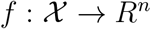 denote a function over input space 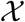 that maps to real-valued scalar outputs. This function can be modeled as a random draw from a GP:

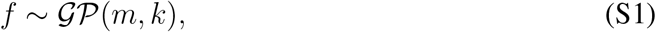

where *m* is a mean function specifying the expected output of the function given input x, and *k* is a kernel (or covariance) function specifying the covariance between outputs:

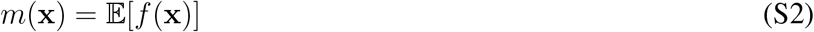

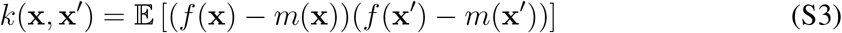

Here, we fix the prior mean to the median value of unscaled payoffs, *m*(x) = 25, and use the kernel function to encode an inductive bias about the expected spatial correlations between rewards (see Radial Basis Function kernel below).

Conditional on observed data 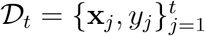, where 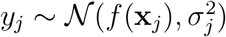 is drawn from the underlying function with added noise 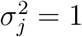, we can calculate the posterior predictive distribution for a new input x_*_ as a Gaussian with mean and variance given by:

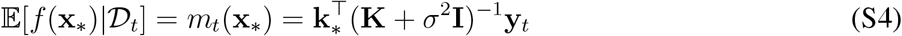

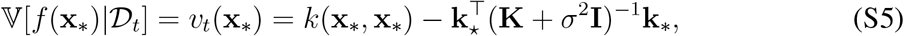

where y = [*y*_1_,…, *y_t_*]^T^, **K** is the *t* × *t* covariance matrix evaluated at each pair of observed inputs, and k* = [*k*(x_1_, x_*_),…, *k*(x_*t*_, x_*_)] is the covariance between each observed input and the new input x_*_.

##### Radial Basis Function kernel

We use the Radial Basis Function (RBF) kernel as a component of the 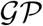 algorithm of generalization. The RBF kernel specifies the correlation between inputs x and x′ as

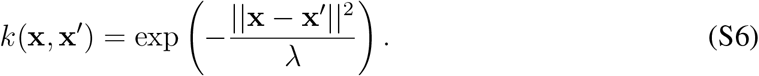

This kernel defines a universal function learning engine based on the principles of Bayesian regression and can theoretically model any stationary function. Note that sometimes the RBF kernel is specified as 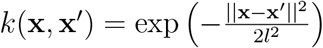 whereas we use λ = 2*l*^2^ as a more psychologically interpretable formulation. Intuitively, the RBF kernel models the correlation between points as an exponentially decreasing function of their distance. Here, λ modifies the rate of correlation decay, with larger λ-values corresponding to slower decays, stronger spatial correlations, and smoother functions. As λ → ∞, the RBF kernel assumes functions approaching linearity, whereas as λ → 0, there ceases to be any spatial correlation, with the implication that learning happens independently for each discrete input without generalization (similar to the assumption of the Mean Tracker model described below). We treat λ as a free parameter, and use cross-validated estimates to make inferences about the extent to which participants generalize.

##### Mean Tracker

The Mean Tracker model is implemented as a Bayesian updating model, which assumes the average reward associated with each option is constant over time (i.e., no temporal dynamics), as is the case in our experiment. In contrast to the GP regression model (which also assumes constant means over time), the Mean Tracker learns the rewards of each option independently, by computing an independent posterior distribution for the mean *μ_j_* for each option *j*. We implemented a version that assumes rewards are normally distributed (as in the GP model), with a known variance but unknown mean, where the prior distribution of the mean is a normal distribution. This implies that the posterior distribution for each mean is also a normal distribution:

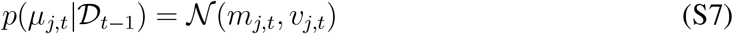

For a given option *j*, the posterior mean *m_j,t_* and variance *v_j,t_* are only updated when it has been selected at trial *t*:

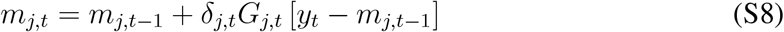

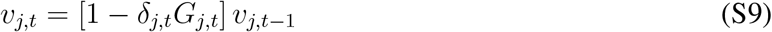

where *δ_j,t_* =1 if option *j* is chosen on trial *t*, and 0 otherwise. Additionally, *y_t_* is the observed reward at trial *t*, and *G_j,t_* is defined as:

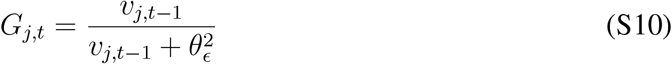

where 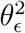 is the error variance, which is estimated as a free parameter. Intuitively, the estimated mean of the chosen option *m_j,t_* is updated based on the difference between the observed value *y_t_* and the prior expected mean *m*_*j,t*-1_, multiplied by *G_j,t_*. At the same time, the estimated variance *v_j,t_* is reduced by a factor of 1 – *G_j,t_*, which is in the range [0,1]. The error variance 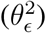 can be interpreted as an inverse sensitivity, where smaller values result in more substantial updates to the mean *m_j,t_*, and larger reductions of uncertainty *v_j,t_*. We set the prior mean to the median value of unscaled payoffs *m*_*j*,0_ = 25 and the prior standard deviation to 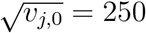.

#### Sampling strategies

Given the normally distributed posteriors of the expected rewards, which have mean *μ*(x) and uncertainty (formalized here as standard deviation) *σ*(x), for each search option x (for the Mean Tracker, we let *μ*(x) = *m_j,t_* and 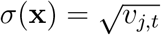, where *j* is the index of the option characterized by x), we assess different sampling strategies that (combined with a softmax choice rule, Eq. S14) make probabilistic predictions about where participants would search next.

##### Upper Confidence Bound sampling

Given the posterior predictive mean *μ*(x) and its attached standard deviation 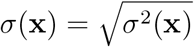, we calculate the upper confidence bound using a weighted sum

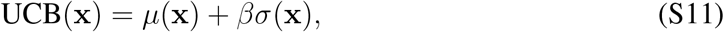

where the exploration factor *β* determines how much reduction of uncertainty is valued (relative to exploiting known high-value options). We estimate *β* as a free parameter indicating participants’ tendency towards directed exploration.

##### Mean Greedy Exploitation

A special case of the Upper Confidence Bound sampling strategy (with *β* = 0) is a greedy exploitation component that only evaluates points based on their expected rewards

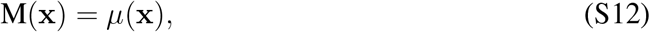

This sampling strategy only samples options with high expected rewards, i.e. greedily exploits the environment.

##### Variance Greedy Exploration

Another special case of the Upper Confidence Bound sampling strategy (with *β* → ∞) is a greedy exploration component which only samples points based on their predictive standard deviation

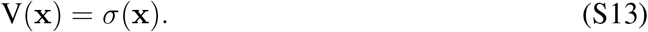

This sampling strategy only cares about reducing uncertainty without attempting to generate high rewards.

### Model comparison

We use maximum likelihood estimation (MLE) for parameter estimation, and cross-validation to measure out-of-sample predictive accuracy. A softmax choice rule transforms each model’s predictions into a probability distribution over options:

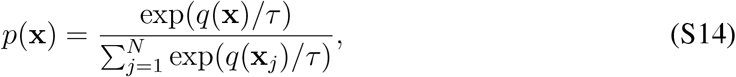

where *q*(x) is the predicted value of each option x for a given model (e.g., *q*(x) = UCB(x) for the UCB model), and *τ* is the temperature parameter. Lower values of *τ* indicate more concentrated probability distributions, corresponding to more precise predictions and therefore less random sampling. All models include *τ* as a free parameter. Additionally, we estimate λ (generalization parameter) for the Gaussian Process regression model and 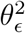 (error variance) for the Mean Tracker model. Finally, we estimate *β* (exploration bonus) for the Upper Confidence Bound sampling strategy.

#### Cross validation

We fit all combinations of models and sampling strategies—per participant—using cross-validated MLE implemented via a Differential Evolution algorithm for optimization. Parameter estimates are constrained to positive values in the range [exp(—5), exp(5)]. We use leave-one-round-out cross-validation to iteratively form a training set by leaving out a single round, computing a MLE on the training set, and then generating out-of-sample predictions on the remaining round. This is repeated for all combinations of training set and test set. This cross-validation procedure yields one set of parameter estimates per round, per participant, and out-of-sample predictions for 200 choices per participant overall (rounds 2-9 × 25 choices).

#### Predictive accuracy

Prediction error (computed as log loss) is summed up over all rounds (apart from the tutorial and the bonus rounds), and is reported as *predictive accuracy*, using a pseudo-*R*^2^ measure that compares the total log loss prediction error for each model to that of a random model:

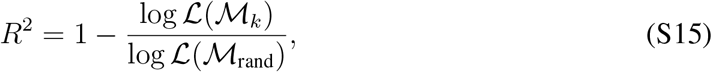

where 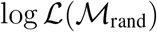 is the log loss of a random model (i.e., picking options with equal probability) and 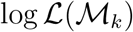 is the log loss of model *k*’s out-of-sample prediction error. Intuitively, *R*^2^ = 0 corresponds to prediction accuracy equivalent to chance, while *R*^2^ = 1 corresponds to theoretical perfect predictive accuracy, since 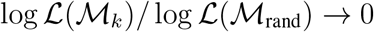 when 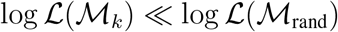.

#### Parameter estimates

All parameter estimates were included in the statistical analyses, although we exclude outliers larger than 5 in Figures 1e, S4, S8, S9, and in Table S1. For GP-UCB estimates, 1.1% of λ-estimates, for 3.4% of *β*-estimates, and 2.7% of *τ*-estimates were removed this way. The presence of these outliers motivated us to use non-parametric tests (without removing outliers) to compare the different parameters across age groups, in order to achieve more robustness. However, the significance of our test results does not change even if we use parametric t-tests, but remove outliers before performing these tests. After removing values higher than 5, we still find that adults show higher λ-estimates than older children (*t*(103) = 3.88, *p* < .001, *d* = 0.76, 95% CI=[0.36,1.16], *BF* > 100), who do not differ in their λ-estimates from younger children (*t*(108) = 1.00, *p* = .32, *d* = 0.19, 95% CI=[−0.19, 0.57], *BF* = 0.3). Moreover, the β-estimates are higher for older children than for adults (*t*(103) = 4.41, *p* < .001, *d* = 0.87, 95% CI=[0.46,1.28], *BF* > 100), but do not differ between the two groups of children *t*(108) = 1.41, *p* = .160, *d* = 0.27, 95% CI=[−0.11, 0.66], *BF* = 0.5). Crucially, the random exploration parameters *τ* do neither differ between older children and adults (*t*(103) = .55, *p* = .58, *d* = 0.11, 95% CI=[−0.28, 0.50], *BF* = 0.2), nor do they differ between the two groups of children (*t*(108) = 1.74, *p* = .08, *d* = 0.33, 95% CI=[—0.72, 0.05], *BF* = 0.8).

Our criterion for outlier removal is less strict than other criteria, such as removing values higher than 1.5×IQR. In our data, this would remove 1.8% of λ-estimates, 5.1% of *β*-estimates, and 11.2% of *τ*-estimates. For completeness, we also reanalyzed the data after performing Tukey’s procedure of outlier removal, where we find the same results. Adults still show a higher λ-estimates than older children (*t*(102) = 3.87, *p* < .001, *d* = 0.76, 95% CI=[0.36,1.17], *BF* > 100), who do not differ from younger children (*t*(108) = 1.04, *p* = .30, *d* = 0.20, 95% CI=[–0.18, 0.58], *BF* = 0.3). The *β*-estimates are again higher for older children than for adults (*t*(100) = 4.69, *p* <.001, *d* = 0.93, 95% CI=[0.51,1.34], *BF* > 100), and do not differ between the two groups of children (*t* (103) = 1.06, *p* = .29, *d* = 0.21, 95% CI=[−0.18,0.60], *BF* = 0.3). Finally, estimates for *τ*-parameter do not differ between adults and older children (*t*(103) = 1.36, *p* = .18, *d* = 0.26, 95% CI=[−0.12, 0.66], *BF* = 0.5), nor between the two groups of children (*t*(108) = 0.96, *p* = .34, *d* = 0.18, 95% CI=[−0.38, 0.38], *BF* = 0.3). Taken together, our results hold no matter which method of outlier removal is applied.

### Full model comparison results

Figure S1 shows the log-scaled Bayes Factor of every combination of each learning model with each sampling strategy, compared in terms of predictive accuracy to every other combination, and separated by the different age groups. Results are based on one-sided Bayesian **t**-tests. We find that the GP-UCB model wins against every other combination, resulting in large Bayes factors overall, but also for every individual age group.

In addition to these Bayes factors for comparisons between individual models, we also compare the cross-validated log-loss at the group level using (predictive) Bayesian model selection, estimating each model’s protected probability of exceedance, which is the likelihood that the proportion of participants generated using a given combination exceeds the proportion of participants generated using all other combinations, while controlling for the chance rate. This analysis revealed that the overall hypothesis of all models performing equally well was rejected by a Bayesian omnibus test at *p* < .001. Moreover, the protected probability of exceedance was virtually 1 for the GP-UCB model, where 1 is the upper theoretical limit. This result is also obtained if breaking down the analysis by the different age groups, again always leading to a protected probability of exceedance of 1 for the GP-UCB model.

Finally, we report how many participants are best predicted by each combination of a model and a sampling strategy. The results of this comparison are shown in Figure S2 (see also Table S1). As before, the GP-UCB model is the best overall model and also predicts most participants best for every age group. Overall, the GP-UCB model predicted 106 out of 160 participants best. In the different age categories, the GP-UCB model best predicted 32 out of 55 younger children, 44 out of 55 older children, and 30 out of 50 adult participants. Therefore, the GP-UCB model was by far the best model in our comparison.

**Figure S1.**
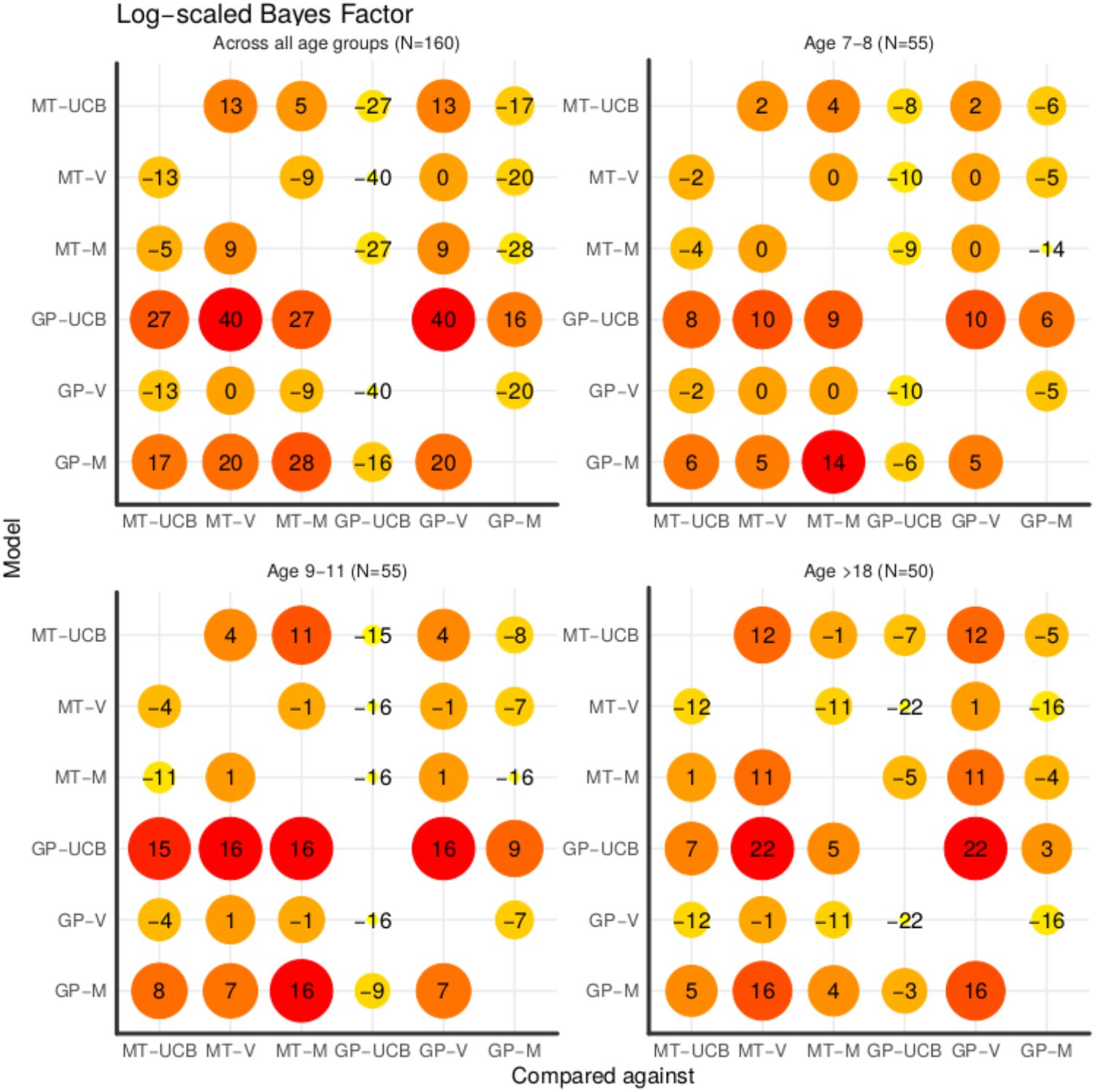
Log-scaled Bayes factor for every pair-wise model comparison, between a model on the y-axis compared against a model on the x-axis. Numbers indicate log_10_ of the Bayes Factor, with a value of 1 indicating that the alternative hypothesis is around 10-times more likely than the null hypothesis, whereas a value of −1 means that the null hypothesis is around 10-times more likely than the alternative hypothesis. Larger Bayes Factors are accompanied by larger circles and darker shades of red. Comparisons show both learning models—the Gaussian Process (GP) and the Mean Tracker (MT)—combined with every sampling strategy, Upper Confidence Bound sampling (UCB), Variance-greedy sampling (V), and Mean-greedy sampling (M).

**Figure S2.**
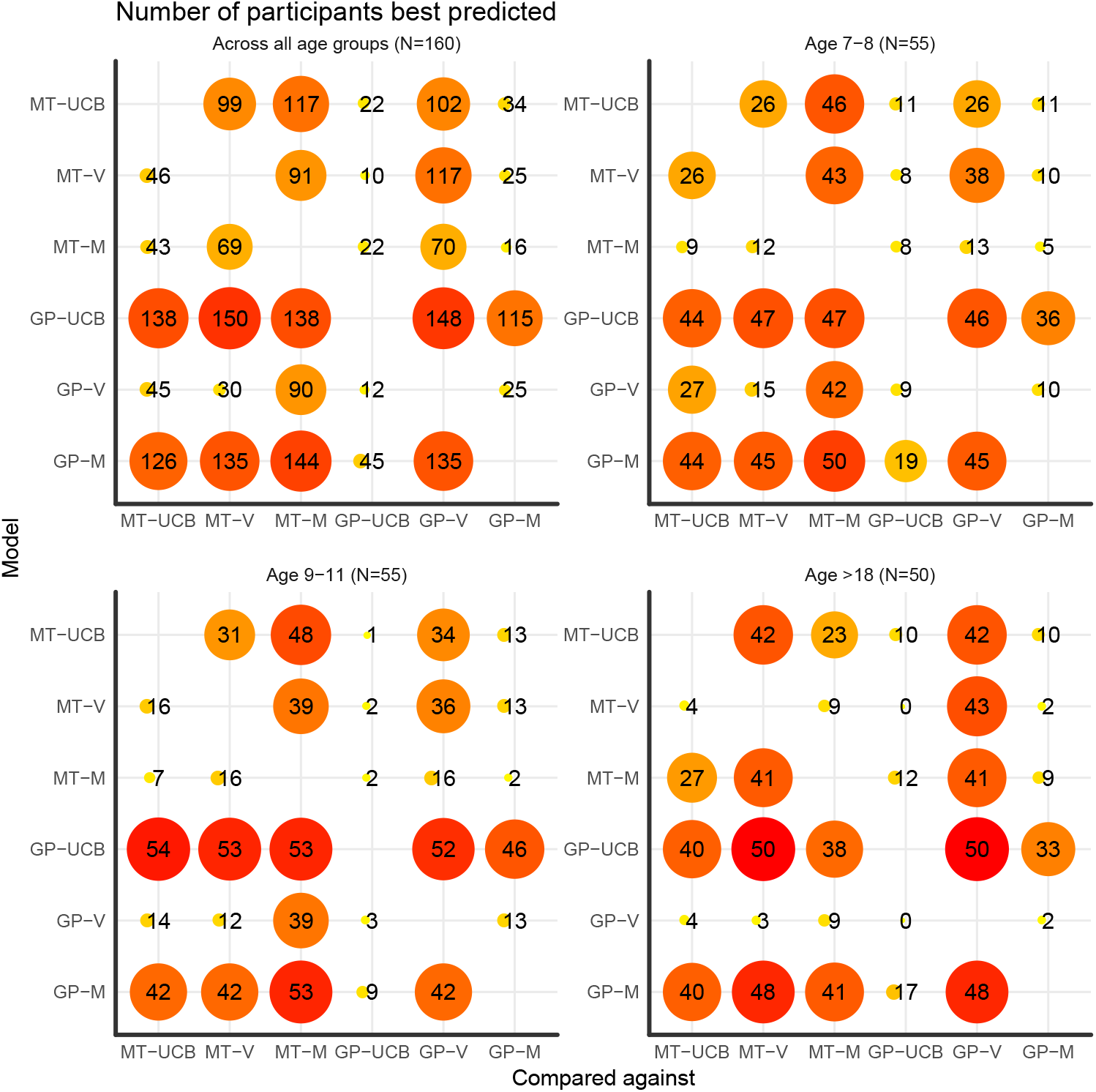
Number of participants best predicted for every pair-wise model comparison, comparing a model on the y-axis with a model on the x-axis. Numbers show absolute counts and can only be interpreted in relation to the overall number of participants per group (shown in brackets at the top of each panel). A larger proportion of participants best predicted are accompanied by larger circles and darker shades of red. Comparisons show both models—the Gaussian Process (GP) and the Mean Tracker (MT)— combined with every sampling strategy, Upper Confidence Bound sampling (UCB), Variance-greedy sampling (V), and Mean-greedy sampling (M).

### Model recovery

We present model recovery results that assess whether or not our predictive model comparison procedure allows us to correctly identify the true underlying model. To assess this, we generated data based on each individual participant’s mean parameter estimates (excluding outliers larger than 5 as before). More specifically, for each participant and round, we use the cross-validated parameter estimates to specify a given model, and then generate new data in the attempt to mimic participant data. We generate data using the Mean Tracker and the GP regression model. In all cases, we use the UCB sampling strategy in conjunction with the specified learning model. We then utilize the same cross-validation method as before in order to determine if we can successfully identify which model has generated the underlying data. Figure S3 shows the cross-validated predictive performance for the simulated data.

#### Recovery Results

**Figure S3.**
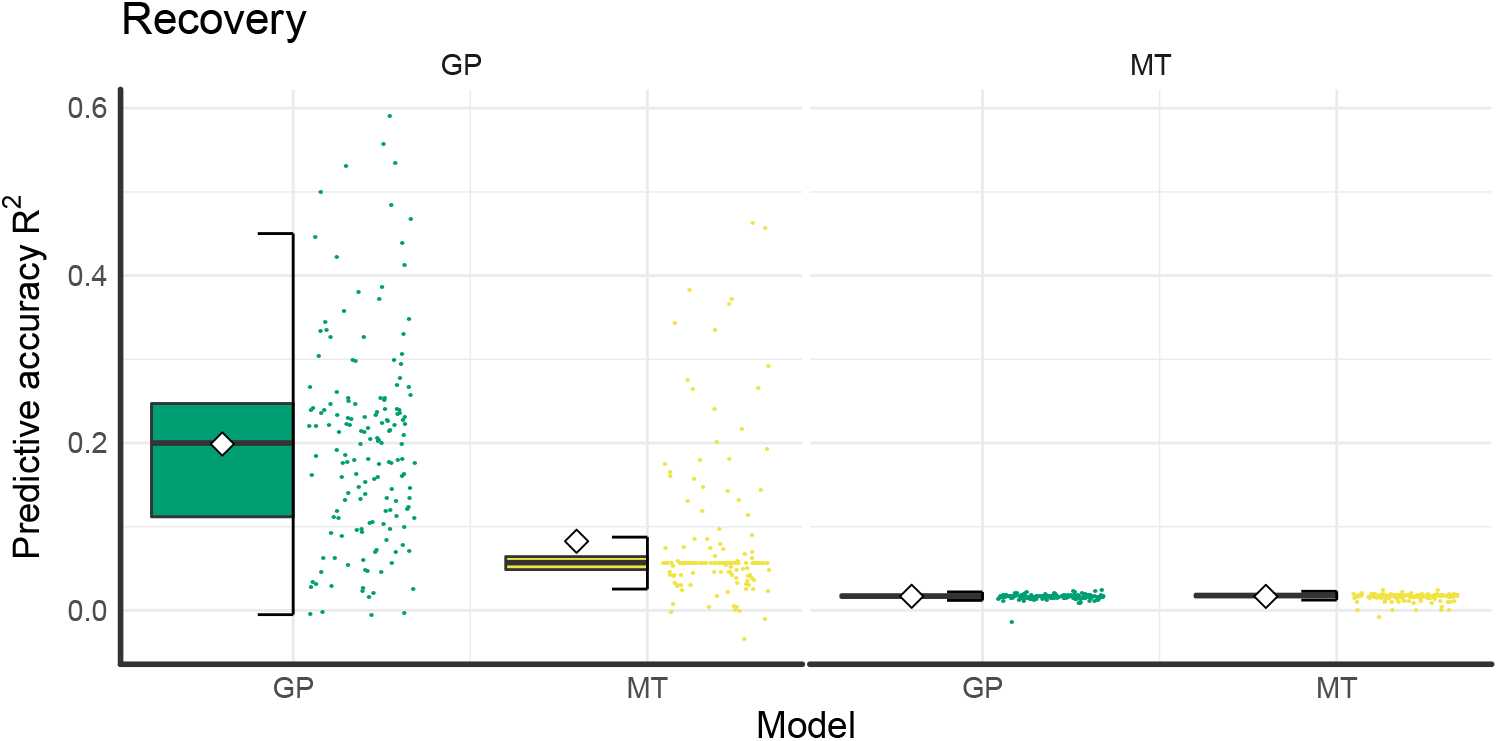
Tukey box plots of model recovery results including individual data points (points) and overall means (diamonds). **Left:** Model performance based on data generated by the GP-UCB model specified with participant parameter estimates. The matching GP-UCB model makes better predictions than a mismatched MT-UCB model. **Right:** Model performance based on data generated by the MT-UCB model specified with participant parameter estimates. Both GP and MT models predict this data poorly, and perform similar to a random model.

Our predictive model comparison procedure shows that the Gaussian Process model is a better predictor for data generated from the same underlying model, whereas the Mean Tracker model is only marginally (if at all) better at predicting data generated from the same underlying model. This suggests that our main model comparison results are robust to Type II errors, and provides evidence that the better predictive accuracy of the GP model on participant data is unlikely due to differences in model mimicry.

When the Mean Tracker model generates data using participant parameter estimates, the same Mean Tracker model performs better than the GP model (*t*(159) = 1.42, *p* = .16, *d* = 0.11, 95% CI=[–0.12,0.33], *BF* = 4.1) and predicts 86 out of 160 simulated participants best. Notice, however, that both models perform poorly in this case, with both models achieving an average pseudo-r-squared of around *R*^2^ = —0.04. This also shows that the Mean Tracker is not a good generative model of human-like behavior in our task.

When the Gaussian Process regression model has generated the underlying data, the same model performs significantly better than the Mean Tracker model (*t*(159) = 18.6, *p* < .001, *d* = 1.47, 95% CI=[1.22,1.72], *BF* > 100) and predicts 153 of the 160 simulated participants best. In general, both models perform better when the GP has generated the data with the GP achieving a predictive accuracy of *R*^2^ = .20 and the MT of achieving *R*^2^ = .12. This makes our finding of the Gaussian Process regression model as the best predictive model even stronger as -technically- the Mean Tracker model can mimic parts of its behavior. Moreover, the resulting values of predictive accuracy are similar to the ones we found using participant data. This indicates that our empirical values of predictive accuracy were as good as possible under the assumption that a GP model generated the data.

### Parameter Recovery

Another important question is whether the reported parameter estimates of the GP-UCB model are reliable and recoverable. We address this question by assessing the recoverability of the three underlying parameters, the length-scale λ, the directed exploration factor *β*, and the random exploration (temperature) parameter *τ* of the softmax choice rule. We use the results from the model recovery simulation described above, and correlate the empirically estimated parameters used to generate data (i.e., the estimates based on participants’ data), with the parameter estimates of the recovering model (i.e., the MLE from the cross-validation procedure on the simulated data). We assess whether the recovered parameter estimates are similar to the parameters that were used to generate the underlying data. We present parameter recovery results for the Gaussian Process regression model using the UCB sampling strategy. We report the results in Figure S4, with the generating parameter estimate on the x-axis and the recovered parameter estimate on the y-axis.

**Figure S4.**
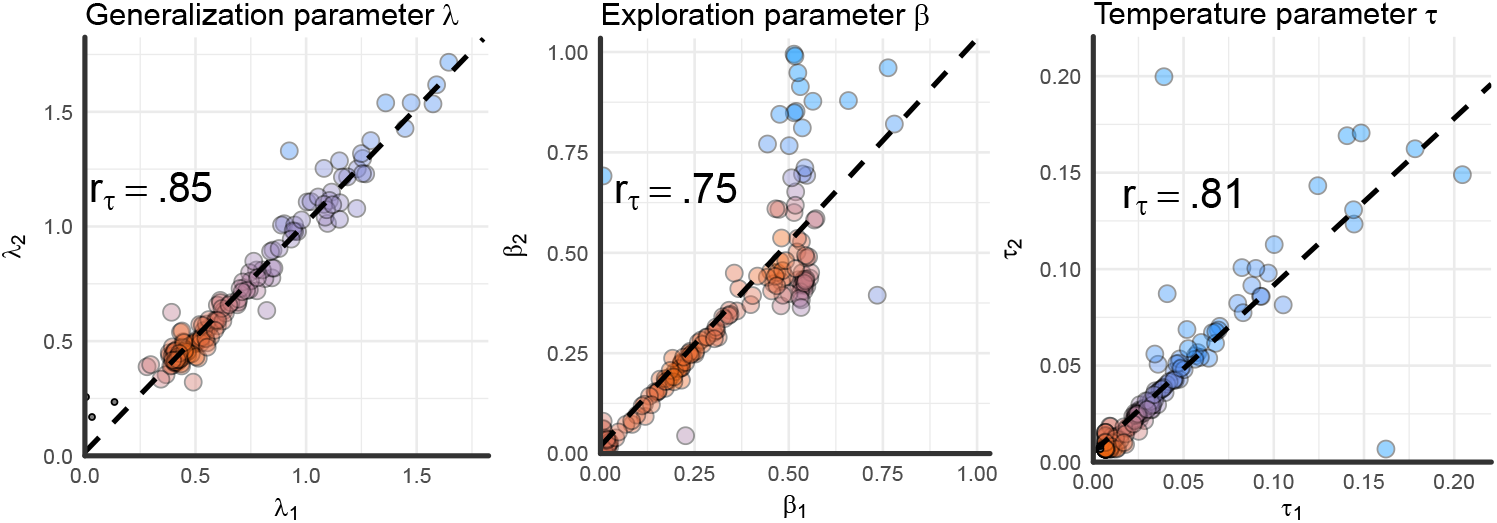
Parameter recovery results. The generating parameter estimate is on the x-axis and the recovered parameter estimate is on the y-axis. The generating parameter estimates are from the cross-validated participant parameter estimates, which were used to simulate data (see Model recovery). Recovered parameter estimates are the result of the cross-validated model comparison (see Model comparison) on the simulated data. While the cross-validation procedure yielded 8-estimates per participant, one for each round, we show the mean estimate per (simulated) participant. The dashed line shows a linear regression on the data, while Kendall’s rank correlation *r_τ_* is shown in the plot. For readability, colors represent the bivariate kernel density estimate, with red indicating higher density.

The correlation between the generating and the recovered length-scale *λ* is *r_τ_* = .85, 95% CI=[0.81,0.90], *p* < .001, the correlation between the generating and the recovered exploration factor *β* is *r_τ_* = 0.75, 95% CI=[0.68, 0.82], *p* < .001, and the correlation between the generating and the recovered softmax temperature parameter *τ* is *r_τ_* = 0.81, 95% CI=[0.76, 0.87], *p* < .001.

These results show that the correlation between the generating and the recovered parameters is very high for all parameters. Thus, we have strong evidence to support the claim that the reported parameter estimates of the GP-UCB model are recoverable, reliable, and therefore interpretable. Importantly, we find that estimates for *β* (exploration bonus) and *τ* (softmax temperature) are indeed recoverable, providing evidence for the existence of a *directed* exploration bonus, as a separate phenomena from *random* exploration in our behavioral data.

Next, we analyze whether or not the same differences between the parameter estimates that we found for the experimental data can also be found for the simulated data. Thus, we compare the recovered parameter estimates from the data generated by the estimated parameters for the different age groups. This comparison shows that the recovered data exhibits the same characteristics as the empirical data. The recovered λ-estimates for simulated data from adults was again larger than the recovered lambda estimates for older children (*U* = 2021, *r_τ_* = 0.33, 95% CI=[0.20, 0.48], *p* < .001, *BF* > 100), whereas there was no difference between the recovered parameters for the two simulated groups of children (*U* = 1800, *p* = .08, *r_τ_* = 0.14, 95% CI=[–.02, 0.29], *BF* = 1). As in the empirical data, the recovered estimates also showed a difference between age groups in their directed exploration behavior such that the recovered *β* was higher for simulated older children than for simulated adults (*U* = 730, *p* < .001, *r_τ_* = 0.33, 95% CI=[0.20, 0.47], *BF* > 100), whereas there was no difference between the two simulated groups of children (*U* = 1730, *p* = .19, *r_τ_* = 0.10, 95% CI=[–0.05,0.26], *BF* = 0.6). There was no difference between the different recovered *τ*-parameters (max-*BF* = 0.5). Thus, our model can also reproduce similar group differences between generalization and directed exploration as found in the empirical data.

#### Counter-factual parameter recovery

Another explanation of the finding that children differ from adult participants in their directed exploration parameter *β* but not in their random exploration parameter *τ* could be that the softmax temperature parameter *τ* can sometimes track more of the random behavioral difference between participants than the directed exploration parameter *β*. If the random exploration parameter *τ* indeed tends to absorb more of the random variance in the data than the directed exploration parameter *β*, then perhaps one is always more likely to find differences in *β* rather than differences in *τ*. To assess this claim, we simulate data using our GP-UCB model as before but swap participants’ parameter estimates of *β* with their estimates of *τ* and vice versa. Ideally, this simulation can reveal whether it is possible for our method to find differences in *τ* but not *β* in a counter-factual parameter recovery where the age groups differ in their random but not their directed exploration behavior. Thus, we generate data from the swapped GP-UCB model and then use our model fitting procedure to assess the GP-UCB model’s parameters from this generated data. The results of this simulation reveal that simulated adults do not differ from simulated older children in their estimated directed exploration parameters *β* (*U* = 1170, *r_τ_* = 0.11, 95% CI=[–0.05, 0.27], *p* = .19, *BF* = 0.7). Furthermore, the two simulated children groups also do not differ in terms of their directed exploration parameter *β* (*U* = 1696, *r_τ_* = 0.09, 95% CI=[–0.07, 0.25], *p* = .27, *BF* = 0.4). However, the random exploration parameter *τ* is estimated to be somewhat higher for simulated older children than for simulated adults (*U* = 1771, *r_τ_* = 0.21, 95% CI=[0.06, 0.36], *p* = .01, *BF* = 2.5) and shows no difference between the two simulated children groups (*U* = 1631, *p* = .48, *r_τ_* = 0.06, 95% CI=[−0.10, 0.21], *BF* = 0.4). This means that the GP-UCB model can pick up on differences in random exploration as well and therefore that our findings are unlikely due to a false positive.

**Figure S5.**
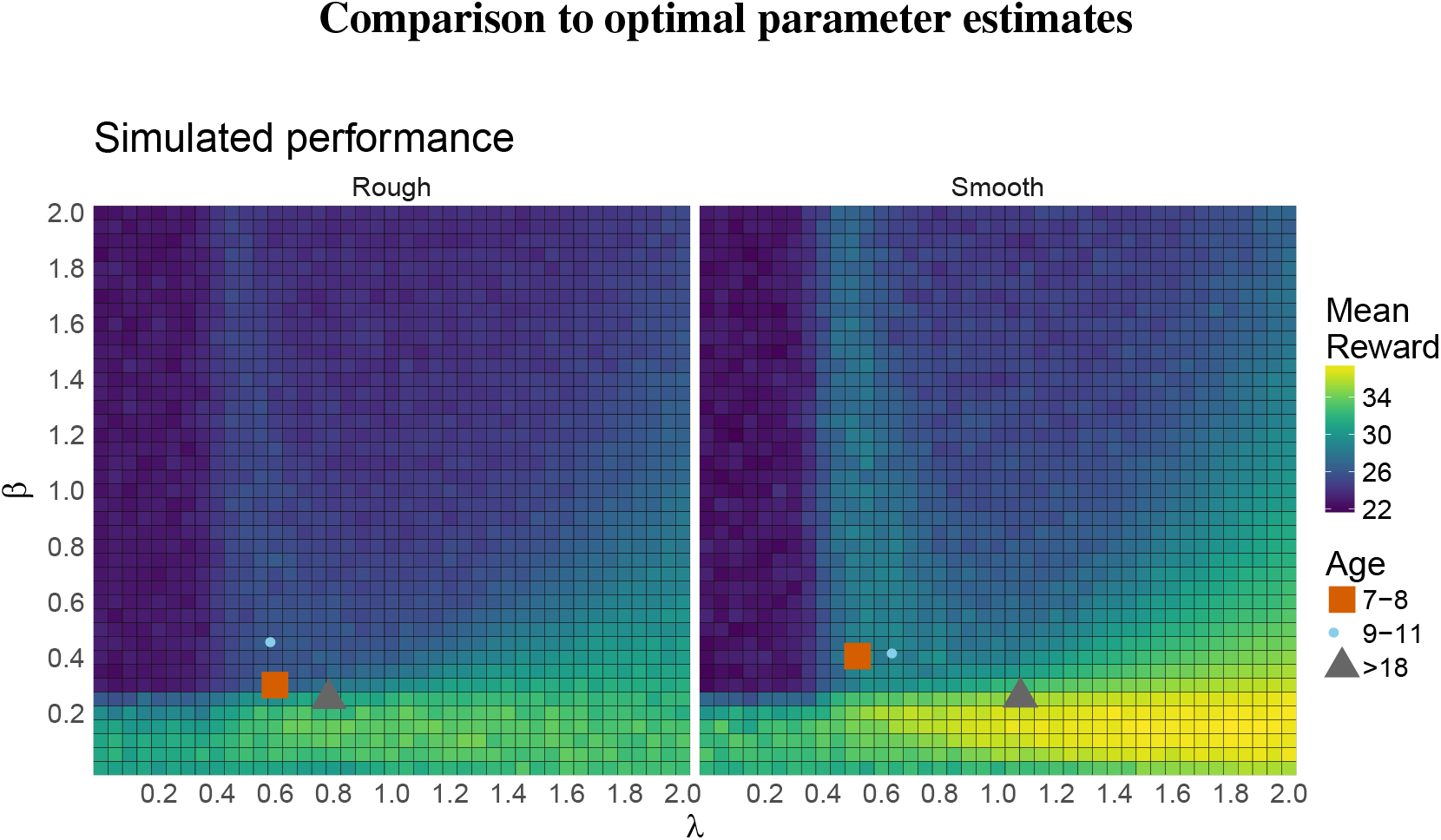
Simulated performance for different parameter values of λ (generalization) and *β* (directed exploration bonus) in the rough (left) and smooth (right) environment. Each tile shows the mean performance over 100 replications on each environment type. The mean participant parameter estimates (separated by age group) are overlaid.

We compare participants’ parameter estimates to optimally-performing estimates of the GP-UCB model (Fig. S5). For this, we simulate the GP-UCB model on both the smooth and the rough environments for the same number of trials as participants experienced, and track performance for each run. Since there were no meaningful differences for the random exploration parameter *τ*, we set *τ* = 0.03 (i.e., the median over all participants) for all simulations. We vary the parameter values of the directed exploration bonus *β* and the generalization parameter λ to all permutations of values stemming from [0.05, 0.1, …, 2], leading to 1,600 differently parameterized models in total. We then run each model for 100 replications on both the smooth and the rough environments individually, always calculating mean performance over all runs. Figure S5 shows the performance of different λ-*β*-combinations with participants’ parameter estimates overlaid.

We extract the best parameters of this simulation by using all parameters that are not significantly different in their performance from the overall best-performing parameters using an α-level of 0.05. The best-performing parameters for the rough condition have a median generalization parameter of λ = 0.95 (range: 0.65-1.30) and a median exploration parameter of *β* = 0.15 (range: 0.1-0.15). The best-performing parameters for the smooth condition have a median generalization parameter of λ = 1.78 (range: 1.4-2) and a median exploration parameter of *β* = 0.15 (range: 0.05-0.2). Unsurprisingly, adults’ parameter estimates are closer to best-performing parameters than children’s parameter estimates. These simulations also replicate earlier findings by Wu et al. (2018) showing that lower values of λ (i.e., undergeneralization) can lead to better performance than values of *λ* that are higher and closer to the true underlying *λ* that generated the environments.

### Further Behavioral Analyses

#### Learning over trials and rounds

We analyze participants learning over trials and rounds using a hierarchical Bayesian regression approach. Formally, we assume the regression weight parameters *θ_x_* for *x* ∈ {0,1,2} are hierarchically distributed as

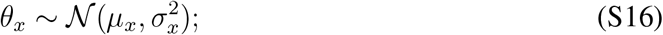

we further assume a weakly informative prior of the means and standard deviation over the regression equation, defined as:

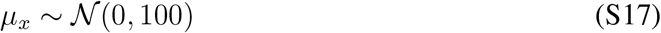

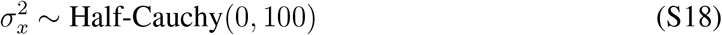

**Figure S6.**
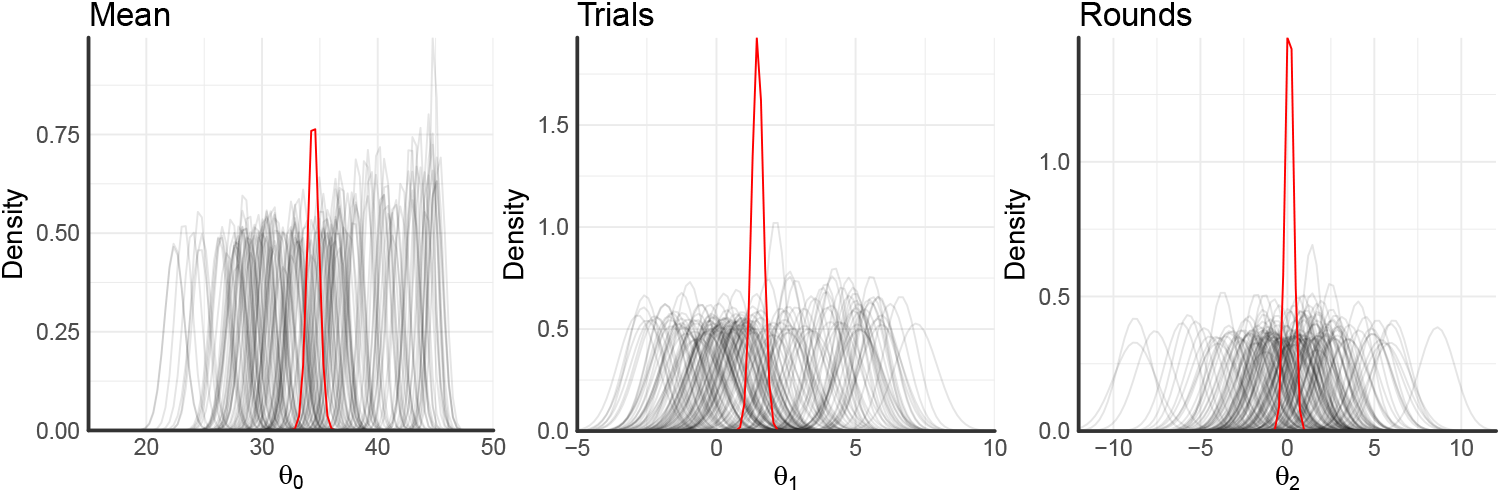
Posterior regression coefficients of mean performance and the effects of both trials and rounds onto participants’ rewards. Individual lines densities correspond to participant-wise estimates, whereas red lines show hierarchical estimates.

We fit one hierarchical model of means and standard deviations over all participants using Hamiltonian Markov chain Monte Carlo sampling as implemented in the `PyMC`-environment. This yields hierarchical estimates for each regression coefficient overall as well as individual estimates for each participant. Doing so for a model containing an intercept (mean performance), a standardized coefficient of the effect of trials on rewards, as well as a standardized coefficient of the effect of rounds on rewards results into the posterior distributions shown in Figure S6.

The overall posterior mean of participants’ rewards is estimated to be 34.4 with a 95%-credible set of [33.5, 35.4]. The standardized effect of trials onto rewards is estimated to be 1.48 with a 95%-credible set of [1.1,1.9], indicating a strong effect of learning over trials. The standardized effect of rounds onto participants’ rewards is estimated to be 0.11 with a 95% credible set of [−0.4,0.6], indicating no effect. Taken together, these results indicate that participants performed well above the chance-level of 25 overall and improved their score greatly over trials. They did not, however, learn or adapt their strategies across rounds.

#### Reaction times

We analyze participants log-reaction times as function of previous reward, age group, and condition. Reaction times are filtered to be smaller than 5000ms and larger than 100ms (3.05% removed in total). We find that reaction times are larger for the rough as compared to the smooth condition (see Fig. S7a; *t*(158) = 4.44, *p* < .001, *d* = 0.70, 95% CI=[0.38,1.02], *BF* > 10). Moreover, adults are somewhat faster than children of age 9-11 (t(103) = —2.60, 95% CI=[0.12, 0.90], *p* = .01, *d* = 0.51, *BF* = 4). The difference in reaction times between children of age 9-11 and children of age 7-8 is only small (*t*(108) = 2.27, *p* = .03, *d* = 0.43, 95% CI=[0.05, 0.81], *BF* = 1.97).

**Figure S7.**
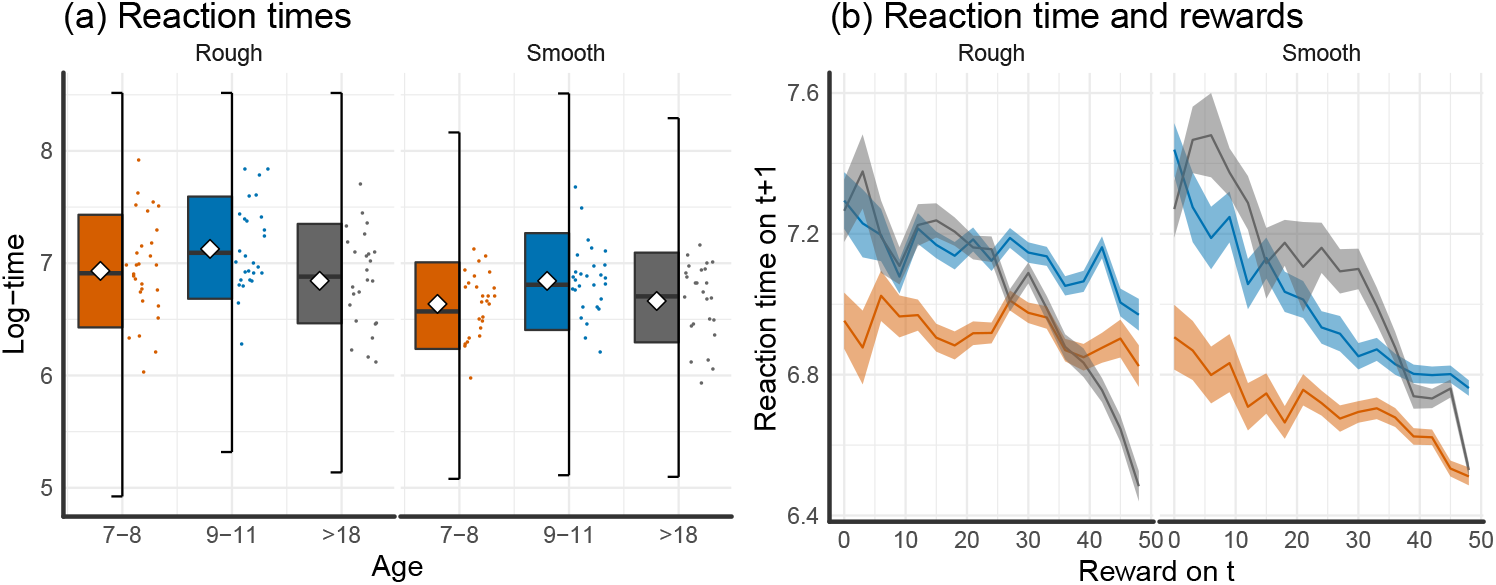
Log-reaction times. (a) Reaction time by age group and condition. (b) Reaction times as a function of previous reward.

We also analyze how much a previously found reward influences participants’ reaction times, i.e. whether participants slow down after a bad outcome and/or speed up after a good outcome (see Fig. S7b). The correlation between the previous reward and reaction times is negative overall with *r* = —0.18, 95% CI=[–0.23, –0.13], *t*(159) = —15, *p* < .001, *BF* > 100, indicating that larger rewards lead to faster reaction times, whereas participants might slow down after having experienced low rewards. This effect is even stronger for the smooth as compared to the rough condition (*t*(158) = –4.43, *p* < .001, *d* = 0.70, 95% CI=[0.38,1.02], *BF* > 100). Moreover, this effect is also stronger for adults than for older children (t(103) = –5.51, *p*< .001, *d* = 1.08, 95% CI=[0.67,1.49], *BF* > 100) and does not differ between the two groups of children (*t*(108) = 1.93, *p* = .06, *d* = 0.37, 95% CI=[–0.01, 0.75], *BF* = 1.05).

**Figure S8.**
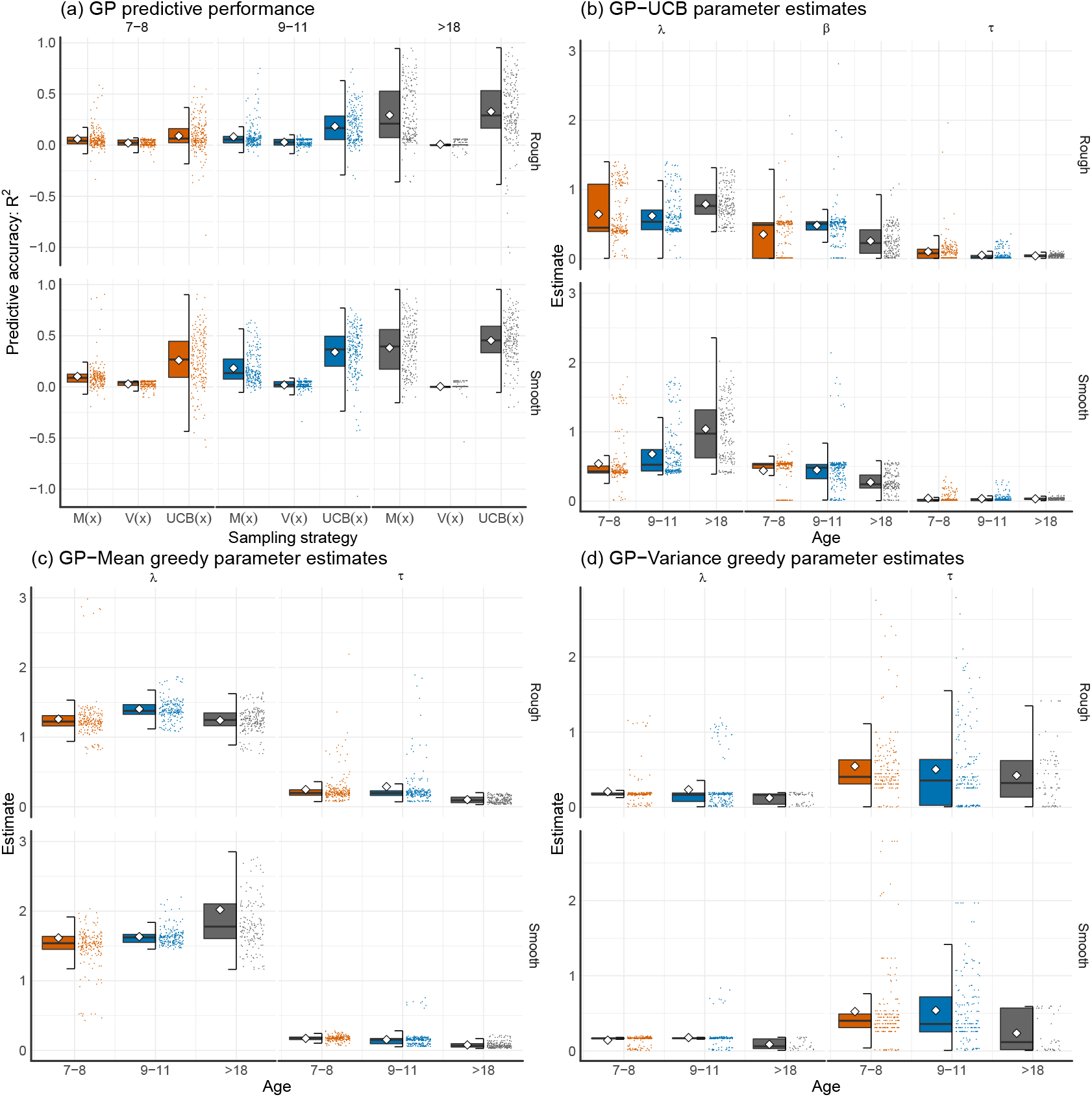
Predictive accuracy (*R*^2^) and parameter estimates for the different Gaussian Process-models by age group and condition. All Tukey box plots include raw data points and mean shown as diamond.

**Figure S9.**
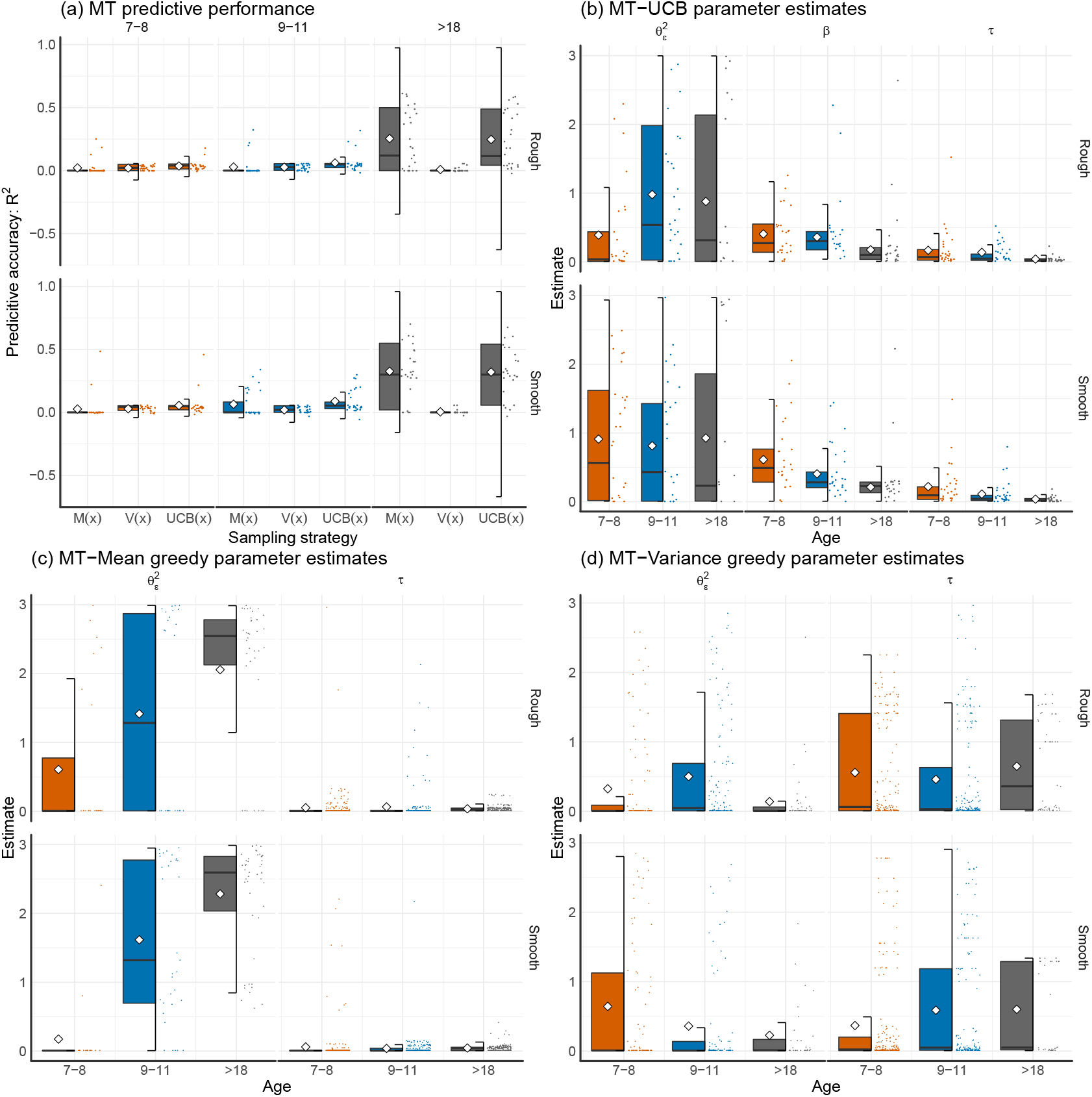
Predictive accuracy (*R*^2^) and parameter estimates for the different Bayesian Mean Tracker-models by age group and condition. All Tukey box plots include raw data points and mean shown as diamond.

**Table S1.** Full modeling results.

| Model | $R^2$ | # Best | Log-loss | Pr. of<br>Exceed. | Generalization<br>$\lambda$ | Exploration<br>$\beta$ | Error Var.<br>$\sqrt{\theta_\epsilon^2}$ | Softmax<br>$\tau$ |
| --- | --- | --- | --- | --- | --- | --- | --- | --- |
| **Overall:** |  |  |  |  |  |  |  |  |
| MT-Mean greedy | 0.02 | 10 | 103.9 | 0 | – | – | 2.98 | 0.01 |
| MT-Variance greedy | 0.01 | 3 | 102.7 | 0 | – | – | 0.58 | 0.08 |
| MT-UCB | 0.05 | 5 | 99.3 | 0 | – | 0.41 | 1.60 | 0.16 |
| GP-Mean greedy | 0.09 | 34 | 94.4 | 0 | 1.46 | – | – | 0.16 |
| GP-Variance greedy | 0.02 | 2 | 102.1 | 0 | 0.18 | – | – | 0.56 |
| GP-UCB | 0.28 | 106 | 74.8 | 1 | 0.59 | 0.45 | – | 0.03 |
| **Age 7-8:** |  |  |  |  |  |  |  |  |
| MT-Mean greedy | 0.00 | 3 | 103.9 | 0 | – | – | 0.53 | 0.01 |
| MT-Variance greedy | 0.00 | 3 | 100.8 | 0 | – | – | 1.22 | 0.08 |
| MT-UCB | 0.03 | 3 | 100.5 | 0 | – | 0.47 | 1.20 | 0.11 |
| GP-Mean greedy | 0.07 | 14 | 96.9 | 0 | 1.37 | – | – | 0.18 |
| GP-Variance greedy | 0.01 | 0 | 102.7 | 0 | 0.17 | – | – | 0.41 |
| GP-UCB | 0.14 | 32 | 89.2 | 1 | 0.46 | 0.48 | – | 0.03 |
| **Age 9-11:** |  |  |  |  |  |  |  |  |
| MT-Mean greedy | 0.01 | 0 | 103.9 | 0 | – | – | 2.86 | 0.01 |
| MT-Variance greedy | 0.01 | 0 | 101.7 | 0 | – | – | 0.65 | 0.08 |
| MT-UCB | 0.05 | 0 | 99.3 | 0 | – | 0.33 | 1.34 | 0.07 |
| GP-Mean greedy | 0.08 | 9 | 95.1 | 0 | 1.52 | – | – | 0.17 |
| GP-Variance greedy | 0.01 | 2 | 101.5 | 0 | 0.17 | – | – | 0.38 |
| GP-UCB | 0.26 | 44 | 76.6 | 1 | 0.74 | 0.53 | – | 0.02 |
| **Adults:** |  |  |  |  |  |  |  |  |
| MT-Mean greedy | 0.29 | 7 | 72.8 | 0 | – | – | 3.38 | 0.03 |
| MT-Variance greedy | 0.00 | 0 | 103.4 | 0 | – | – | 0.13 | 0.87 |
| MT-UCB | 0.29 | 2 | 74.2 | 0 | – | 0.18 | 2.94 | 0.03 |
| GP-Mean greedy | 0.33 | 11 | 69.1 | 0 | 1.38 | – | – | 0.07 |
| GP-Variance greedy | 0.01 | 0 | 103.9 | 0 | 0.10 | – | – | 0.23 |
| GP-UCB | 0.40 | 30 | 62.7 | 1 | 0.85 | 0.24 | – | 0.03 |

*Note:* Columns indicate (from left to right) the average predictive accuracy of each model ( $R^2$ ), how many participants each model best predicted (# Best), the average log-loss over all 8 rounds of predictions (Log-loss), the protected probability of exceedance (Pr. of Exceed.), the GP's generalization parameter ( $\lambda$ ), the directed exploration parameter ( $\beta$ ) for all models involving a UCB sampling strategy, the MT's error variance ( $\sqrt{\theta_\epsilon^2}$ ), and the random exploration (softmax temperature) parameter ( $\tau$ ). Parameter estimates are based on median over per-participant mean parameter estimates (excluding estimates larger than 5 as outliers).

## Footnotes

1 We also assessed if there was a correlation between age and parameter estimates for the adult participants. This revealed no relation between age and λ (*r* = −0.11, *t*(48) = −0.73, *p* = .47, *BF* = 0.4), *β* (*r* = 0.15, *t*(48) = −1.03, *p* = .31, *BF* = 0.5) or *τ* (*r* = −0.09, *t*(48) = −0.62, *p* = .53, *BF* = 0.4). However, these results should be interpreted with caution as they are only based on 50 subjects. Future research should try to further map out the developmental trajectories of these parameters across the whole lifespan.

2 Note that interpreting estimates of inferior computational models can be problematic and should only be done with caution.

## Notes

#### Summary of Updates

Final psych science version.

## References

Auer, P. (2002). Using confidence bounds for exploitation-exploration trade-offs. Journal of Machine Learning Research, 3, 397–422.

Bellman, R. (1952). On the theory of dynamic programming. Proceedings of the National Academy of Sciences, 38, 716–719.

Blanco, N. J., Love, B. C., Ramscar, M., Otto, A. R., Smayda, K., & Maddox, W. T. (2016). Exploratory decision-making as a function of lifelong experience, not cognitive decline. Journal of Experimental Psychology: General, 145, 284–297.

Bonawitz, E. B., van Schijndel, T. J., Friel, D., & Schulz, L. (2012). Children balance theories and evidence in exploration, explanation, and learning. Cognitive Psychology, 64, 215–234.

Cauffman, E., Shulman, E. P., Steinberg, L., Claus, E., Banich, M. T., Graham, S., & Woolard, J. (2010). Age differences in affective decision making as indexed by performance on the iowa gambling task. Developmental Psychology, 46, 193–207.

Davidson, D. (1991). Developmental differences in children’s search of predecisional information. Journal of Experimental Child Psychology, 52, 239–255.

Davidson, D. (1996). The effects of decision characteristics on children’s selective search of predecisional information. Acta Psychologica, 92, 263–281.

Decker, J. H., Otto, A. R., Daw, N. D., & Hartley, C. A. (2016). From creatures of habit to goal-directed learners: Tracking the developmental emergence of model-based reinforcement learning. Psychological Science, 27, 848–858.

Frank, M. J., Doll, B. B., Oas-Terpstra, J., & Moreno, F. (2009). Prefrontal and striatal dopaminergic genes predict individual differences in exploration and exploitation. Nature Neuroscience, 12, 1062–1068.

Gershman, S. J. (2018). Deconstructing the human algorithms for exploration. Cognition, 173, 34–42.

Gopnik, A., Griffiths, T. L., & Lucas, C. G. (2015). When younger learners can be better (or at least more open-minded) than older ones. Current Directions in Psychological Science, 24, 87–92.

Gopnik, A., O’Grady, S., Lucas, C. G., Griffiths, T. L., Wente, A., Bridgers, S.,… Dahl, R. E. (2017). Changes in cognitive flexibility and hypothesis search across human life history from childhood to adolescence to adulthood. Proceedings of the National Academy of Sciences, 114, 7892–7899.

Hagen, J. W., & Hale, G. A. (1973). The development of attention in children. ETS Research Report Series, 1973.

Hartley, C. A., & Somerville, L. H. (2015). The neuroscience of adolescent decision-making. Current Opinion in Behavioral Sciences, 5, 108–115.

Josef, A. K., Richter, D., Samanez-Larkin, G. R., Wagner, G. G., Hertwig, R., & Mata, R. (2016). Stability and change in risk-taking propensity across the adult life span. Journal of Personality and Social Psychology, 111, 430.

Keresztes, A., Bender, A., Bodammer, N., Lindenberger, U., Shing, Y. L., & Werkle-Bergner, M. (2017). Hippocampal maturity promotes memory distinctiveness in childhood and adolescence. Proceedings of the National Academy of Sciences, 114, 9212–9217.

Klahr, D. (1982). Nonmonotone assessment of monotone development: An information processing analysis. In S. Strauss & R. Stavy (Eds.), U-shaped behavioral growth (pp. 63–86). New York: Academic Press.

Lucas, C. G., Bridgers, S., Griffiths, T. L., & Gopnik, A. (2014). When children are better (or at least more open-minded) learners than adults: Developmental differences in learning the forms of causal relationships. Cognition, 131, 284–299.

Lucas, C. G., Griffiths, T. L., Williams, J. J., & Kalish, M. L. (2015). A rational model of function learning. Psychonomic Bulletin & Review, 22, 1193–1215.

Mata, R., Wilke, A., & Czienskowski, U. (2013). Foraging across the life span: is there a reduction in exploration with aging? Frontiers in Neuroscience, 7, 53.

Mehlhorn, K., Newell, B. R., Todd, P. M., Lee, M. D., Morgan, K., Braithwaite, V. A.,… Gonzalez, C. (2015). Unpacking the exploration–exploitation tradeoff: A synthesis of human and animal literatures. Decision, 2, 191–215.

Palminteri, S., Kilford, E. J., Coricelli, G., & Blakemore, S.-J. (2016). The computational development of reinforcement learning during adolescence. PLoS Computational Biology, 12, e1004953.

Piaget, J. (1964). Part i: Cognitive development in children: Piaget development and learning. Journal of Research in Science Teaching, 2, 176–186.

Rasmussen, C., & Williams, C. (2006). Gaussian Processes for machine learning. Cambridge, MA, USA: MIT Press.

Riedmiller, M., Hafner, R., Lampe, T., Neunert, M., Degrave, J., Van de Wiele, T.,… Springenberg, J. T. (2018). Learning by playing-solving sparse reward tasks from scratch. arXiv preprint arXiv:1802.10567.

Rouder, J. N., Speckman, P. L., Sun, D., Morey, R. D., & Iverson, G. (2009). Bayesian t tests for accepting and rejecting the null hypothesis. Psychonomic Bulletin & Review, 16, 225–237.

Ruggeri, A., & Lombrozo, T. (2015). Children adapt their questions to achieve efficient search. Cognition, 143, 203–216.

Schulz, E., Konstantinidis, E., & Speekenbrink, M. (2017). Putting bandits into context: How function learning supports decision making. Journal of Experimental Psychology: Learning, Memory, and Cognition, 44, 927–943.

Schulz, L. E. (2015). Infants explore the unexpected. Science, 348, 42–43.

Shepard, R. N. (1987). Toward a universal law of generalization for psychological science. Science, 237, 1317–1323.

Sollich, P. (1999). Learning curves for Gaussian processes. In Advances in Neural Information Processing Systems (pp. 344–350).

Somerville, L. H., Sasse, S. F., Garrad, M. C., Drysdale, A. T., Abi Akar, N., Insel, C., & Wilson, R. C. (2017). Charting the expansion of strategic exploratory behavior during adolescence. Journal of Experimental Psychology: General, 146, 155–164.

Srinivas, N., Krause, A., Kakade, S. M., & Seeger, M. (2009). Gaussian process optimization in the bandit setting: No regret and experimental design. arXiv preprint arXiv:0912.3995.

Turing, A. (1950). Computing intelligence and machinery. Mind, 59, 433–460.

Tymula, A., Belmaker, L. A. R., Roy, A. K., Ruderman, L., Manson, K., Glimcher, P. W., & Levy, I. (2012). Adolescents’ risk-taking behavior is driven by tolerance to ambiguity. Proceedings of the National Academy of Sciences, 109, 17135–17140.

van Doorn, J., Ly, A., Marsman, M., & Wagenmakers, E.-J. (2017). Bayesian latent-normal inference for the rank sum test, the signed rank test, and Spearman’s p. arXiv preprint arXiv:1712.06941.

White, J. M. (2013). The role of delayed consequences in human decision-making. (Unpublished doctoral dissertation). Princeton University.

Wilson, R. C., Geana, A., White, J. M., Ludvig, E. A., & Cohen, J. D. (2014). Humans use directed and random exploration to solve the explore–exploit dilemma. Journal of Experimental Psychology: General, 143, 2074–2081.

Wu, C. M., Schulz, E., Speekenbrink, M., Nelson, J. D., & Meder, B. (2018). Generalization guides human exploration in vast decision spaces. Nature Human Behaviour, 2, 915–924.

